# Hyppo-X: A Scalable Exploratory Framework for Analyzing Complex Phenomics Data

**DOI:** 10.1101/159954

**Authors:** Methun Kamruzzaman, Ananth Kalyanaraman, Bala Krishnamoorthy, Stefan Hey, Patrick S. Schnable

## Abstract

Phenomics is an emerging branch of modern biology that uses high throughput phenotyping tools to capture multiple environmental and phenotypic traits, often at massive spatial and temporal scales. The resulting high dimensional data represent a treasure trove of information for providing an in-depth understanding of how multiple factors interact and contribute to the overall growth and behavior of different genotypes. However, computational tools that can parse through such complex data and aid in extracting plausible hypotheses are currently lacking. In this paper, we present `Hyppo-X`, a new algorithmic approach to visually explore complex phenomics data and in the process characterize the role of environment on phenotypic traits. We model the problem as one of unsupervised structure discovery, and use emerging principles from algebraic topology and graph theory for discovering higher-order structures of complex phenomics data. We present an open source software which has interactive visualization capabilities to facilitate data navigation and hypothesis formulation. We test and evaluate `Hyppo-X` on two real-world plant (maize) data sets. Our results demonstrate the ability of our approach to delineate divergent subpopulation-level behavior. Notably, our approach shows how environmental factors could influence phenotypic behavior, and how that effect varies across different genotypes and different time scales. To the best of our knowledge, this effort provides one of the first approaches to systematically formalize the problem of hypothesis extraction for phenomics data. Considering the infancy of the phenomics field, tools that help users explore complex data and extract plausible hypotheses in a data-guided manner will be critical to future advancements in the use of such data.

## 1 Introduction

High-throughput technologies are beginning to change the way we observe and measure the natural world. In medicine, physicians are using imaging and other specialized sampling devices to keep a longitudinal log of patients’ drug/therapy response and other disease-related phenotypes. In agricultural biotechnology, phenotyping technologies such as cameras and LiDARs are being used to measure physiological and morphological features of crops in fields. Further, advancements in genotyping technologies (sequencing) have made it possible to characterize and track genetic diversity and changes at a high resolution, and decode genetic markers that are key to performance traits. Taken together, advancements in these technologies are leading to a rapid explosion of high-dimensional data, obtained from a variety of sources.

A distinctive feature of these inherently highdimensional data sets is that their generation is motivated more based on the availability and easy access to high-throughput technology as opposed to specific working hypotheses. While there are some broad high-level questions or research themes that motivate the collection of data, the specific questions that relate to testable hypothesis and eventual discoveries (e.g., what genetic variations impact a physical trait, or how a combination of environmental variables control a phenotype) are *not* readily available *a priori*.

Consider the case of plant phenomics [1], [2]. Understanding how different crop varieties or *genotypes* (*G*) interact with *environments* (*E*) to produce different varying performance traits (*phenotypes* (*P*)) is a fundamental goal of modern biology (*G × E* → *P*) [3], [4]. To address this fundamental albeit broad goal, plant biologists and farmers have started to widely deploy an array of high-throughput sensing technologies that measure tens of crop phenotypic traits in the field (e.g., crop height, growth characteristics, photosynthetic activity). These technologies, comprising mostly of camera and other recording devices, generate a wealth of images (visual, infrared, thermal) and time-lapse videos that represent a detailed set of observations of a crop population as it develops over the course of the growing season. Additionally, environmental sensors help in collecting daily field measurements that represent the growth conditions. Furthermore, through the use of sophisticated genotyping technologies, the genotypes of the different crop varieties are also cataloged.

From this medley of plant genotypes, phenotypes, and environmental measurements, scientists aim to extract plausible hypotheses that can be field-tested and validated. However, the task remains significantly challenging, mainly due to the dearth of automated software capabilities that are capable of handling complex, high-dimensional data sets. Scatter plots (such as the example shown in Fig. 1) and correlation studies can reveal only high-level correlations and behavioral patterns/differences within data. However, it is common knowledge that different individuals or subgroups of individuals behave differently under similar stimuli. For instance, while it is useful to know that a given environmental variable (e.g., humidity) shows an overall positive correlation to a performance trait (say, crop height), such high-level correlations obfuscate the variations within a population—e.g., how different subgroups or genotypes respond to different intervals in the environmental values; or how one environmental variable interacts/interplays with another to affect the performance trait; or how the same genotype expresses variability in its performance under different environments (plasticity).

**Fig. 1.**
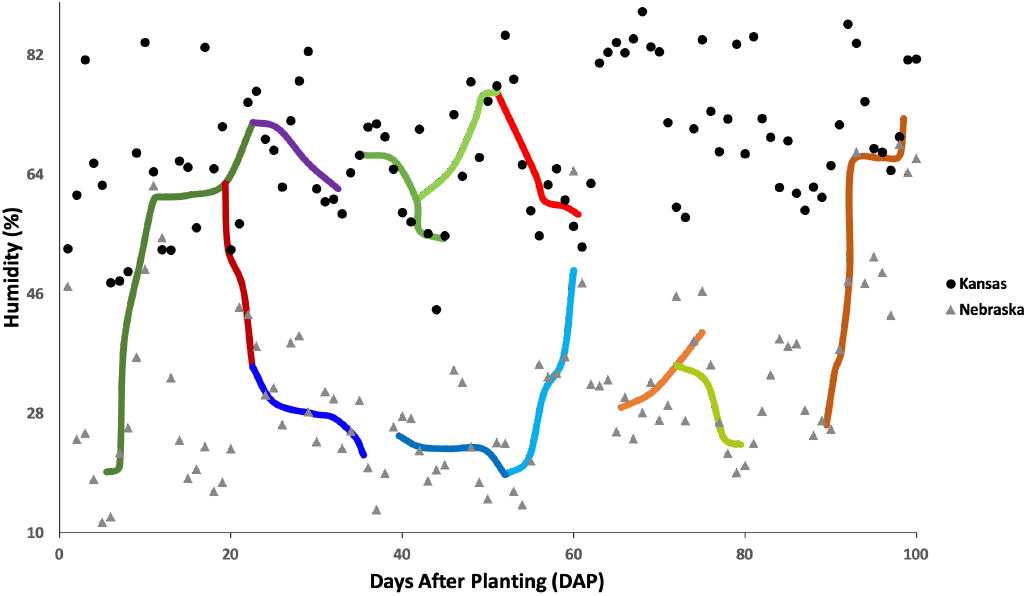
Scatter plot of a maize crop data set containing points grown in two locations—Kansas (KS) and Nebraska (NE). Each data “point” is an [individual, date/time] combination, with x-axis representing the Days After Planting (time) and y-axis representing humidity as an environmental variable. While the scatter plot shows higher humidity values in KS than in NE in general, it does not in itself have the capability to show intra-population variation with respect to a performance variable. To this end, the “interesting paths” generated by our TDA framework can be useful. These paths are shown overlaid on the scatter plot; each path highlights a subset of points that are connected by consistent performance behavior (growth rate in this example). This could help us identify behaviorally-coherent subpopulations within large populations that are otherwise nontrivial to observe.

### 1.1 Our Contributions

In this paper, we present a novel computational approach for extracting hypotheses from high-dimensional data sets such as ones collected in phenomics. We formulate the problem of hypothesis extraction as one of: (a) identifying the key connected structural features of the given data, and (b) exploring the structural features in a way to facilitate extraction of plausible hypotheses.

#### 1.1.1 Structure Identification

Our approach uses emerging principles from *algebraic topology* as the basis to observe and discern structural features from raw phenomics data. Algebraic topology is the field of mathematics dealing with the shape and connectivity of spaces [5], [6]. There are multiple important properties of topology that make it particularly effective for extracting structural features from large, high-dimensional data sets. First, topology studies shapes in a *coordinate-free way*, which enables comparison among data sets from diverse sources or coordinate systems. Second, topological constructions are *not sensitive to small changes in data*, and robust against noise. Third, topology works with *compressed representations* of spaces in the form of *simplicial complexes* (or triangulations) [5], which preserve information relevant to how points are connected. Compared to more traditional techniques such as principal component analysis, multidimensional scaling, and cluster analysis, topological methods are known to be more sensitive to both large and small scale patterns [7].

#### 1.1.2 Topological Object Exploration

While topological representations offer a compact way to represent and explore the data, the problem of how to navigate such representations in order to glean hypothesis information is still unexplored. In this paper, we formulate this problem formally as identifying a) *interesting flares* and b) *interesting paths*. The features we target encapsulate different properties of the data, as detailed below.

**Interesting flares:** Flares show how a subset of points (i.e., subpopulation) branches into smaller subpopulations when exposed to certain environmental stimuli—e.g., a set of plant individuals (or varieties) that shows divergent behavior in their growth characteristics when one of the environmental parameters (say, temperature) crosses a certain value. Identifying such flares could help us identify subpopulations of interest and track their behavioral evolution at a finer granularity of the population.

**Interesting paths:** A *path*, on the other hand, highlights a trail of point clusters along which a “performance” variable increases (or decreases). In other words, a path can reveal different subpopulations that are prevalent in different performance intervals. This can in turn help us contrast different population subtypes or subsets by their performance under different environmental conditions. An illustrative example highlighting this feature is shown in Fig. 1.

We first define these features formally, and then present algorithms to extract them from the topological objects constructed. For ranking purposes, we define a notion of interestingness.

#### 1.1.3 Software

We have implemented our approach as a software tool, which we call `Hyppo-X` (stands for: <u>Hyp</u>othesis extraction for <u>p</u>hen<u>o</u>mics). The tool is available as open source in the GitHub repository [8].

Even though we demonstrate its utility in the context of plant phenomics, our approach can be applied more broadly to other similar application contexts where the goal is to identify interesting subpopulations in general in an unsupervised manner from complex, high-dimensional biological data sets.

## 2 Related work

We are not aware of any other automated or semi-automated hypothesis extraction approaches for high-dimensional data sets. In what follows, we present some related work, both in topology and in plant phenomics, in order to put our contributions in context.

### 2.1 Topology and Applications

There are several important properties that make algebraic topology particularly effective for gleaning structural features out of high-dimensional data. First, topology studies shapes in a *coordinate-free way*, which enables comparison among data sets from diverse sources or coordinate systems. Second, topological constructions are *not sensitive to small changes in data*, and robust against noise. Third, topology works with *compressed representations* of spaces in the form of *simplicial complexes* (or triangulations) [5], which preserve information relevant to how points are connected. Compared to more traditional techniques such as principal component analysis, multidimensional scaling, manifold learning, and cluster analysis, topological methods are known to be more sensitive to both large and small scale patterns [7].

Topological data analysis (TDA) has been applied to a wide range of application domains, albeit for mostly visualization purposes [9], [10], [11], [12], [13], [14]. The foundational work in TDA most relevant to this paper was done by Carlsson and coworkers [7]. In [15], they describe a framework called *Mapper* to model and visualize high-dimensional data. Most of this work has been on the visualization front. A topology-based approach was also rated as the best overall entry at an expression QTL (eQTL) visualization competition organized by the BioVis community [16].

### 2.2 Tools for Plant Phenomics

Tools to decode associations between genotypes and phenotypes have been under development for over two decades. These tools look at the genetic variation observed at one or more loci across the genome and study their correlation to quantitative traits. The techniques used can be summarized as follows: i) Linkage mapping usually begins with prior knowledge of the order of genetic markers and the goal of the mapping is to identify which markers co-segregate with a phenotype in a segregating population. It is usually used for traits controlled by fewer genes; ii) Quantitative Trait Locus (QTL) mapping that extends linkage to an interval of co-located markers along the genome; and iii) GWAS is typically used for traits controlled by many genes. Typically all individuals within a diversity panel are scored for both genotypes at many markers AND phenotypes. Statistical approaches are then used to identify statistically significant associations between markers and variation in trait values. In relation to capturing environmental variability, efforts have been sparse. [17] presented an experimental framework supplemented by GWAS to model environmental effects on phenotypes. [18] provided a generalized linear model-based method to capture gene to environment interactions. In another related work, Yang *et al*. [19] study the effect of environmental variables on photosynthesis efficiency in plants using a curve fitting approach.

The approach presented in *this* paper complements the above body of works in several ways including a new way to formulate the problem as one of unsupervised structure discovery, and in its method and capabilities (e.g., compact representation, visualization, and exploratory data analysis).

## 3 Hyppo-X: Our implementation of the Mapper framework

The first step in our approach is to construct topological representations using the connectivity properties of the data. The motivation is to obtain higher order structural information about the high-dimensional data prior to gleaning hypotheses. We present an implementation for the abstract Mapper algorithmic framework [15] for this purpose. In what follows, we describe the details of our implementations. Fig. 2 is a schematic illustration of our approach.

**Fig. 2.**
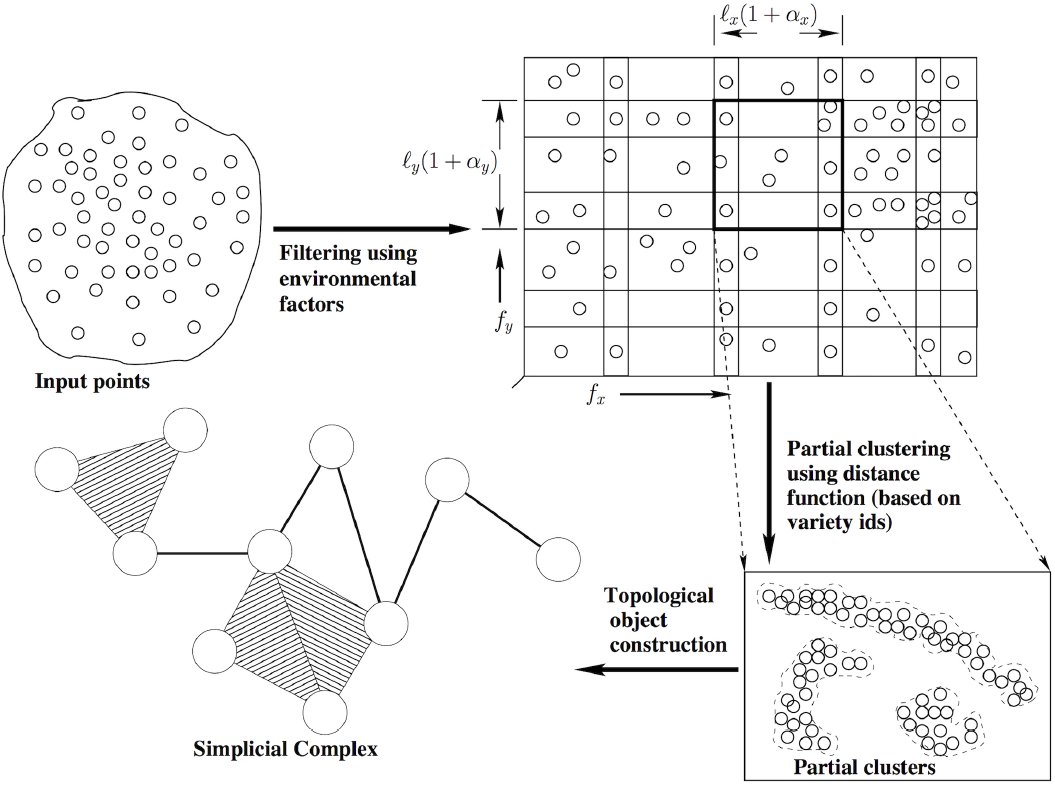
`Hyppo-X` framework for analyzing phenomic data.

**Input:** We are given a set of *n* points 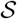 in a *d*-dimensional space, representing the space of interest *X*. In the case of phenomics, a *point* 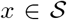 represents a crop individual that is measured at a particular time *t*, and the dimensions represent the attributes which describe the point at that time. These include a set *E* of *m* factors (e.g., time, temperature, humidity, etc.), and a performance trait, the phenotype *p* (e.g., plant height or growth rate). Note that these dimensions represent continuous variables (A point may also have other non-continuous or static variables (e.g., the genotype). For the purpose of our topological representations we will use only the continuous variables).

**Output:** We aim to create a highly compact coordinate-free representation of *X* as a *simplicial complex*, using a clustering (overlapping) of the points in *X* (represented by *P* here).

**Simplicial complex:** A *simplicial complex* is a collection of simplices (nodes, edges, triangles, tetrahedra, etc.) that fit together nicely—all subsimplices of each simplex are included in the collection, and any two simplices that intersect do so in a lower dimensional subsimplex. Specifically, each cluster is represented by a node (0-simplex). Whenever two clusters have a non-empty intersection, we add an edge (1-simplex), and when three clusters intersect, we add a triangle (2-simplex), and so on.

We now provide the main algorithmic details of the approach.

### 3.1 Filtering

The first component of the framework is a continuous function *f: X → Z* to a real-valued parameter space *Z*, called the *filter function*. For each factor *Z_j_*, we define a filter function *f_j_: X → Z_j_*. We generate the open cover 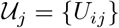 of *Z_j_* as follows:

1. We divide each factor *Z_j_* into *n_j_* intervals (“subregions”), each of length *ℓ_j_*. Thus the entire *d*-dimensional region is divided into *n*_1_ × *n*_2_ × ⋯ × *n_m_* subregions, where each subregion represents a hyper-rectangle of area *ℓ*_1_ × *ℓ*_2_ × ⋯ × *ℓ_m_*. Let the center of *i^th^* hyper-rectangle be {*C*_1*i*_, *C*_2*i*_,…, *C_mi_*}.
2. We fix the center of each hyper-rectangle, and increase the length along each factor *Z_j_* by a certain percentage *α_j_* such that an overlapping region is created between consecutive pairs of the open sets *U_ij_* and *U*_*i*+1,*j*_, i.e., 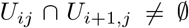. After increasing the length of all sides in this fashion, the new area of the hyper-rectangle is *ℓ*_1_(1 + *α*_1_) × ⋯ × *ℓ_m_*(1 + *α_m_*) (See Section 3.5 for further explanation of how we choose the *α_i_* values). A 2D example is shown in Fig. 2.

We formulate the efficient determination of individual point sets belonging to each hyper-rectangle as a problem of range querying. Specifically, we implement the following querying function:

**Range Query:** *Given X and a hyper-rectangle h, return the subset of points in X that lie in h*.

To run this query efficiently, we use *k*-dimensional hyper-octtrees [20], [21], which is a well known spatial data structure that uses recursive bisection to index a spatially distributed set of points. The compressed version of an *n*-leaves hyper-octree can be constructed in *O*(*n* log *n*) time [20]. Once constructed, a balanced binary search tree that uses the order of the leaves is constructed. Using this auxiliary data structure, in combination with the hyper-octree, enables an *O*(log *n*) worst case search time for both point and cell searches [20]. To answer the regional query for a hyper-rectangle *h*, we perform a top-down traversal of the hyper-octree by selectively retaining only those paths that can include at least one point within *h*. This can be achieved by keeping track of the corners of the cell defined by each internal node in the tree. This approach ensures that each such query can be answered in time that is bounded by the number of points in the hyper-rectangle.

### 3.2 Generation of Partial Clusters

Each open set (hyper-rectangle) computed by applying the filter functions is processed independently for generation of partial clusters. The goal of clustering is to partition the set of points in each hyper-rectangle based on their phenotypic performance.

Let *U* represent an open set of points {*x*_1_, *x*_2_,… *x_t_*}. Note that each point *x* ∈ *U* has a phenotypic trait value denoted by *p*(*x*). We define a *distance function d* based on the phenotypic values of points in *U* as follows. Given two points with trait values *p*(*x_i_*) and *p*(*x_j_*), the distance *d*(*i,j*) = |*p*(*x_i_*) − *p*(*x_j_*)|.

Given *U* and distance function *d*, a *partial clustering* is defined by a partitioning of the points in *U*. We denote the set of partial clusters resulting from any given open set *U* as 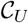. Subsequently, we denote the set of all partial clusters formed from *all* open sets (hyper-rectangles) by 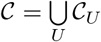.

For the purpose of clustering, any distance-based clustering method can be applied. We implemented a density-based clustering approach very similar to that of DBSCAN [22]. It covers two key points: a) the set of partial clusters generated from within a hyper-rectangle represents a partitioning of those points; and b) two partial clusters generated from a pair of adjacent (overlapping) hyper-rectangles could potentially have a non-empty intersection in points. In fact it is this intersection that renders connectivity among the partial clusters generated, the information for which will be used in the subsequent step of simplicial complex generation.

### 3.3 Construction of Simplicial Complexes

From the set of partial clusters 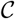, we construct a simplicial complex *M* as follows. We describe the details for the 2D case, where no more than four open sets (hyper-rectangles) can mutually intersect. The extension to higher dimensions is straightforward. Starting with an empty simplicial complex, we implement the following steps:

1. A 0-simplex (or vertex) is added to the simplicial complex *M* for every partial cluster.
2. Next, a 1-simplex (edge) is added to *M* for every nonempty 2-way intersection between any two partial clusters. Note that such intersections could exist only between partial clusters originating from different open sets.
3. Following the same procedure as above, we also add 2-simplices (triangles) and 3-simplices (tetrahedra) to *M* by enumerating only those 3-way and 4-way intersections, respectively, that could be non-empty.

The required multi-way intersections are computed using the range querying function described earlier (in Section 3.1).

The Mapper algorithm [15] produces highly compressed visual representations of high-dimensional data that reveal significant structural aspects. For example, consider the instance where *X* is a set of points in ℝ^2^ sampled from a noisy unit circle (see Fig. 3). We use the height of the points (i.e., their *y*-coordinate values) as the filter function. We consider a cover of *Z*, which is almost [−1,1], into *r* = 3 overlapping intervals, with adjacent intervals overlapping roughly by a third (i.e., *g* = 33%). The pullback cover of *X* then has four pieces, with the subset of points with height in the middle interval forming two connected components. We then use the Euclidean distance between the points (in ℝ^2^) as the distance function to cluster the points in each component using, e.g., single linkage clustering. Thus we get one node per component, which we color from blue to red according to the mean height of the points in each node. We also get four connecting edges capturing the overlap of the clusters. Note that we go from around 20 points in *X* to just four nodes and four edges in the Mapper. At the same time, this highly compact representation captures the underlying structure of *X*—the circle.

**Fig. 3.**
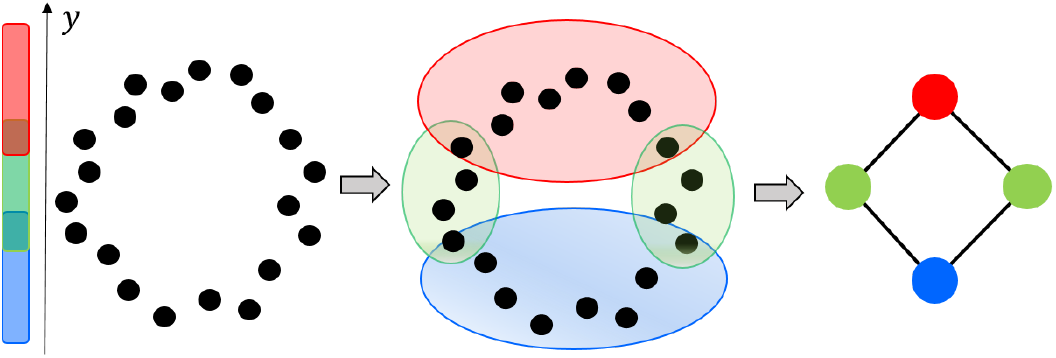
The Mapper algorithm applied to a set of points sampled from a noisy circle. We use the height of the points (*y*-coordinate) as the filter function. We consider a cover of *Z* ≈ [−1,1] using *r* = 3 overlapping intervals, with adjacent intervals overlapping roughly by a third (i.e., *g* = 33%). The final Mapper is shown on the right.

### 3.4 Graph Formulation

We construct a weighted directed graph *G* = (*V, E*) representation of the 1-skeleton of *M* along with some additional information. We set *V* as the set of vertices (0-simplices) of *M*, and *E* as the set of edges (1-simplices) of *M*. We assign directions and weights to the edges as follows. Each vertex *u* ∈ *V* denotes a subset of points from *X* that constitute a partial cluster. We denote this subset as *X*(*u*). We let *g*(*u*) and *f_i_*(*u*) denote the average values of the clustering function *g* (dependent variable) and the filter function *f_i_*, respectively, for all points in *u*:

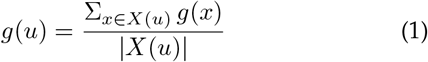

and

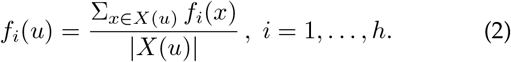

For an edge *e* = (*u, v*) in *E*, we assign as its weight as: *ω*(*e*) = |*g*(*u*) − *g*(*v*)|. Notice *ω*(*e*) ≥ 0 for all edges *e* in *G*. In addition, the direction of the edge e is set from the lower weight vertex to the higher weight vertex—i.e., if *ω*(*u*) ≤ *ω*(*v*) then *e*: *u* → *v*, and *e: v → u* otherwise. We let *n* = |*V*| and *m* = |*E*| denote the numbers of vertices and edges in *G*, respectively.

#### 3.4.1 Edge

If the simplicial complex was constructed using *h* out of the *m* continuous variables (as filter functions), then along each edge, each continuous variable *f_i_* can independently increase or decrease. Since we are trying to link the change of each of these variables relative to the change in phenotype (along an edge), we record a *h*-bit *signature* for each edge.

We assign a h-bit binary signature Sig(*e*) = *b*_1_*b*_2_…*b_h_* to the oriented edge *e* = (*u, v*) (i.e., *e: u → v*) to capture the covariation of *g* and the filter functions *f_i_*. We set *b_i_* = 1 if *f_i_*(*u*) ≤ *f_i_*(*v*), and *b_i_* =0 otherwise.

In other words, let an edge’s direction be *u → v*. Then, if the mean value for the continuous variable *f_i_* increases from *u* to *v*, then the corresponding signature bit is 1; and 0 otherwise.

Fig. 4 illustrates a directed, signed edge in our representation.

**Fig. 4.**
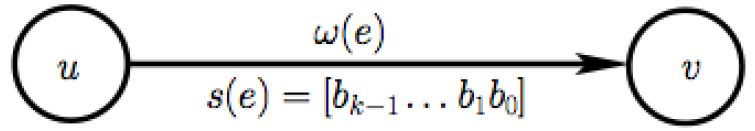
An edge *e* between two intersecting partial clusters (nodes *u* and *v*). The direction of the edge indicates the direction in which the mean phenotypic/performance value increases. The signature *s*(*e*) is a *k*-bit vector that captures the directions of change for each of the *k* filter functions (e.g., environmental variables) along the edge—0 implies decreasing and 1 implies increasing. The *i^th^* bit corresponds to the *i^th^* filter function.

Note that based on the above edge definition, there cannot be any cycles in *G*(*V, E*), making it a Directed Acyclic Graph (DAG).

### 3.5 Persistent homology

We employ the concept of persistent homology [23] to choose the final topological object for further analysis. In particular, the method in which overlapping intervals are chosen (by specifying growing overlap percentages *α_i_*, see Section 3.1) is already guided by this principle. Termed *multiscale mapper*, growing the intervals in this fashion ensures the topological objects formed (at each set of growing *α_i_* values) satisfy a monotonic inclusion property [24]. Hence results from persistent homology could be used to guarantee (theoretical) stability of the topological object formed (in the sense of persistence). At the same time, no implementation of multiscale mapper is known. Instead, we increase each *α_i_* in steps of 2.5%, and construct the topological objects for each set of *α_i_* values. We then construct the persistence barcodes (in dimensions 0, 1, and 2) using the sequence of topological objects formed by employing JavaPlex, a standard software tool for this purpose. We then pick *α_i_*’s such that all three barcodes do not change for values at or higher than the chosen cutoff, ensuring the corresponding topological object chosen is indeed stable.

## 4 Extracting interesting features

In this section, we define two features—flares and paths— that can be extracted from the topological representations we construct (in Section 3), and subsequently describe our algorithms to extract those features. Interesting flares and paths hold two different types of information: flares are more useful to identify subpopulation divergence, whereas paths are more useful to identify and analyze subpopulations over the performance spectrum.

### 4.1 Interesting Flares

We propose a framework to detect and use “flares” (defined below) that characterize branching phenomena in phenomics data sets.

We first construct a directed graph G based on the process discussed in Section 3.4. Given an edge *e* = {*u, v*}, we direct the edge by default from the cluster showing a lower phenotypic performance as measured by its mean phenotypic value to one with the higher mean phenotypic value (see Fig. 4). This scheme allows us to track a trail of clusters that show an improving trajectory in performance by a user-selected phenotypic trait (e.g., yield or plant height). In order to capture branching phenomena effectively, we modify this directing procedure by using mean phenotypic values of *subset of individuals belonging to shared genotypes* between nodes *u* and *v*.

#### Definition 4.1.

A *source* (*terminal*) node in a directed graph is one that has no incoming (outgoing, respectively) edges. A *branching* node in a directed graph is one that has at least two outgoing edges.

Note that, by the above definitions, a source node can also be potentially a branching node. Furthermore, we use the term *simple path* to refer to a path in the graph in which no node, with the possible exception of the sentinel nodes (beginning and ending) of the path, is a branching node.

We define a *stem* and a *branch* associated with a branching node as follows (see Figure 5 for an illustration).

**Fig. 5.**
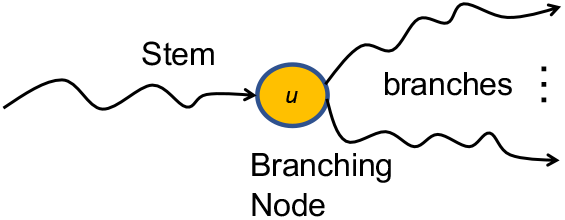
An illustration of a flare.

#### Definition 4.2.

Given a branching node *u*, a *stem* is a possibly empty simple path that ends in *u*.

Note that there can be multiple stems ending at a branching node *u*. There are two classes of such stems—those that are entirely non-overlapping (i.e., simple paths ending at *u* that are otherwise node-disjoint) and those that are nested (i.e., they originate from different starting nodes in the same parent simple path ending at *u*).

#### Definition 4.3.

Given a branching node *u*, a *branch* refers to a non-empty path (simple or not) that originates at *u*.

Note that two branches originating at the same branching node can possibly intersect. Furthermore, there are at least two branches originating at a branching node (by definition of a branching node).

Let *B*(*u*) denote the set of all branches originating at a branching node *u* and *S*(*u*) denote the set of nonoverlapping (i.e., non-nested) stems ending at *u*.

#### Definition 4.4.

We define *a flare* to be a unique combination of a branching node *u*, a stem *s* ∈ *S*(*u*), and a subset *B*′(*u*) ⊆ *B*(*u*). Here, we do *not* enforce that a stem be nonempty, to allow detection of flares strictly originating at a given branching node. However, we *do* enforce that each branch selected is non-empty (i.e., has at least one edge) and that the subset selected *B*′(*u*) ⊆ *B*(*u*) contains at least two or more branches (as illustrated in Fig. 5).

The selection of the stem and branches to include in a flare is computed deterministically as a function of the branching node. Intuitively, the idea is to examine the set of individuals “covered” by the branching node, and then “cast a net” in either direction, on all simple paths leading up to *u* (candidate stems) and on all the branches originating at *u*, as far as there is a non-empty intersection with the individual set of the branching node (see Fig. 6).

**Fig. 6.**
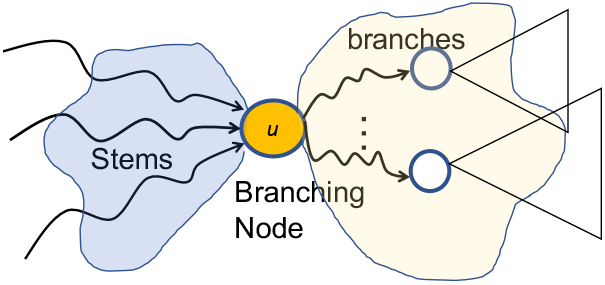
Conceptual illustration of how flares are constructed from a given branching node *u*. Stems are selected from the set of incoming simple paths, and branches are selected from the DAGs rooted at *u*. The boundaries of the selection are determined by “casting a net” on either side of *u* and including all “areas” where there is shared individual coverage.See text above for further details.

The rationale for this selection scheme is as follows. In an application such as phenomics, each “point” included in a cluster is typically a given plant crop (“individual”) observed in certain time and space. Therefore, by the way we construct our topological object using intersections between adjacent clusters, the same individual may continue to appear in a sequence of clusters (i.e., in a path) on either side of a branching node. Therefore, by considering the set of individuals covered by a branching node, and examining how that set distributes itself across the branches, we can discover interesting subpopulation-level variations (or differences in the way they respond to various environmental filters). In a population where there is also a large genetic diversity, one can adapt the same procedure to include the set of genotypes covered (instead of plant individuals).

**Detection of flares:** More formally, let *N*(*u*) denote the set of individuals covered in the cluster corresponding to *u*. Then, we follow the trail of clusters in either direction to incrementally grow the corresponding stem or branch, as follows. For stem computation, we enumerate all the simple paths ending at *u*, and for each such simple path (candidate stem), we begin at the node *v* which is the immediate predecessor of *u* and compute *N*(*v*) ∩ *N*(*u*). If the intersection is non-empty then we include *v* in the current stem and iteratively walk to the next predecessor (until either the simple path terminates or the intersection becomes empty). Note that at each step, we compute the intersection with *N*(*u*).

A similar procedure is carried out to enumerate all branches originating at *u*, walking forward instead of backward, with the caveat that we do not need to restrict the elongation process to only simple paths in the forward direction. In other words, if we encounter another branching node, the algorithm proceeds recursively, except that at every subsequent step going forward from the second branching node, the intersection is computed only relative to the original branching node *u*.

Note that the above procedure is deterministic, in that given a branching node, the reach of a flare involving that branching node is determined by the reach of the set of individuals in *u* on either side of *u* in the DAG. In fact, this procedure would also detect *all* the flares involving *u*. More precisely, the cross-product of *S*(*u*) and *B*′(*u*) (as specified in Definition 4.4) yields the set of all flares involving *u*.

**Scoring flares:** In order to compare and relatively rank flares, we devise a simple scheme to score each flare. Given a flare *f*, we compute its “interestingness score” as follows.

First, we associate a weight to all edges. The weight of an edge is given by the absolute difference in the phenotypic performance (cluster means) between the two corresponding clusters. Intuitively, the larger the performance variation, the more interesting that edge is to a branch. Note that since we use the absolute value of the difference, all edge weights are positive.

We score the flare using its edge weights as follows (see Fig. 7). Note that there is a unique subgraph induced by each flare and that subgraph also will be acyclic (as it is derived from a DAG). Therefore, we perform a simple bottom-up/post-order traversal of that induced DAG, starting at each terminal node and climbing up the parent and the ancestor levels. At each step, we perform a simple gather-scatter way to propagate the scores across levels. More specifically, at a node *u*, all the scores of its child branches are added (“gather”), and the value is then equally divided (“scatter”) among its predecessor branches. The algorithm terminates when it reaches the main branching node *u* of this flare. Once scored, the flares can be rank ordered in the decreasing order of score and displayed.

**Fig. 7.**
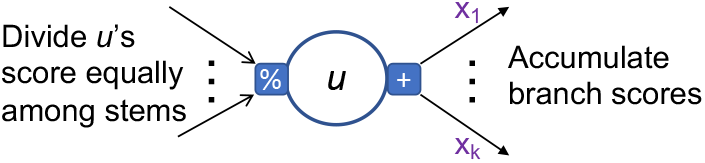
Illustration of how the interestingness score propagates through a flare. Computation proceeds as an accumulation process from the branches to the stems.

In our current implementation, only branches contribute to the score of a flare at a branching node. Stems do not contribute, the rationale being that examining the branches typically suffices for explaining how a population, covered at the branching node, diverges. However, the procedure can be extended to include stem scores as needed. The information contained in the stem is still useful during our subsequent analysis and interpretation.

Our procedure for scoring flares takes running time linear in the size of the flare.

### 4.2 Interesting Paths

We propose a framework to detect and use “paths” (defined below) that helps to identify interesting subpopulation in phenomics data sets.

#### Definition 4.5.

An *interesting k-path* for a given *k* with 1 ≤ *k* ≤ *n* − 1 is a directed path *P* = [*e*_*i*_1__,… *e_i_k__*] of *k* edges in *G*, such that Sig(*e_r_*) is identical for all *r* = *i*_1_,…,*i_k_*. An *interesting path* is a path of arbitrary length in the interval [1, *n* − 1].

#### Definition 4.6.

Given an interesting *k*-path *P* = [*e*_*i*_1__,…,*e_i_k__*] in *G* as specified in Definition 4.5, we define its *interestingness score* as follows.

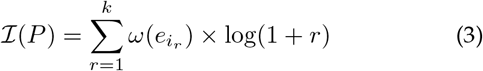

In particular, the contribution of an edge *e* ∈ *P* to 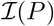 is set to *ω*(*e*) × log(1 + rank(*e, P*)), where rank(*e, P*) is the rank or order of edge *e* as it appears in *P*.

Intuitively, we use the rank of an edge as an inflation factor for its weight—the later an edge appears in the path, the more its weight will count toward the interestingness of the path. This logic incentivizes the growth of long paths. The log function, on the other hand, helps temper this growth in terms of number of edges.

**Optimization Problems:** We now present multiple optimization problems with the broader goal of identifying interesting path(s) that maximize interestingness score(s).

**Max-IP:** Find an interesting path *P* in *G* such that 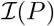 is maximized.

**IP:** Find a collection 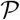 of interesting paths in *G* such that the total interestingness score 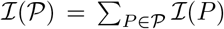 is maximized (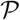 will exactly cover *E*, i.e., each *e ∈ E* is part of exactly one 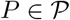).

A detailed analysis the above optimization problems (and related variants) with their respective complexity results and proofs are provided in a separate manuscript [25]. In what follows, we present an exact algorithm for the MAX-IP problem. We also present an efficient heuristic for the IP problem. Both these algorithms are implemented in our `Hyppo-X` framework.

#### 4.2.1 *The* Max-IP *Problem*

The goal of MAX-IP is to identify an interesting path with the maximum interestingness score. We show MAX-IP is P on directed acyclic graphs (DAGs).

##### Lemma 4.7.

MAX-IP *on a directed acyclic graph G* = (*V, E*) *is in P*.

*Proof*. We present a polynomial time algorithm for MAX-IP on a DAG (as proof of Lemma 4.7). The input is a DAG *G* = (*V, E*) with *n* vertices and *m* edges, with edge weights *ω*(*e*) ≥ 0 and signatures Sig(*e*) for all *e ∈ E*. The output is an interesting path *P*\* which has the maximum interestingness score in *G*. We use dynamic programming, with the forward phase computing 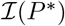 and the backtracking procedure reconstructing a corresponding *P*\*.

Let *T*(*i,j*) denote the score of a maximum interesting path of length *j* edges ending at edge *e_i_* for *i* ∈ [1, *m*]. Since an interesting path could be of length at most (*n* − 1), we have *j* ∈ [1, *n* − 1]. Therefore the values in the recurrence can be maintained in a 2-dimensional table of size *m* × (*n* − 1), as illustrated in Fig. 8. The algorithm has three steps:

- **Initialization:** *T*(*i*, 1) = *ω*(*e_i_*) × log(2), where 1 ≤ *i* ≤ *m*.
- **Recurrence:** For an edge *e* = (*u, v*) ∈ *E*, we define a *predecessor edge* of *e* as any edge *e*′ ∈ *E* of the form *e*′ = (*w, u*) and Sig(*e*′) = Sig(*e*). Let Pred(*e*) denote the set of all predecessor edges of e. Note that Pred(*e*) can be possibly empty. We define the recurrence for *T*(*i,j*) as follows.

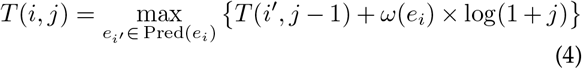
- **Output:** We report the score that is maximum in the entire table. A corresponding optimal path *P*\* can be obtained by backtracking from that cell to the first column.

**Fig. 8.**
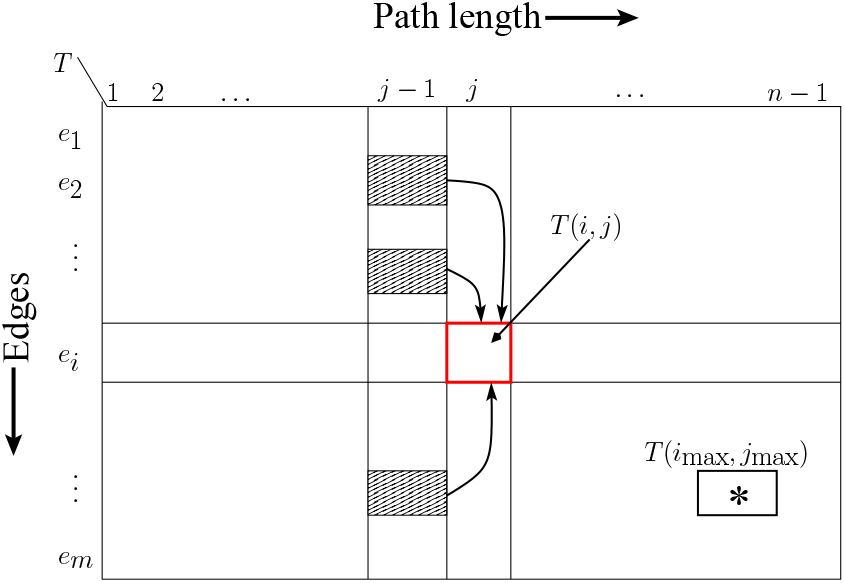
Table *T*(*i,j*) for the Max-IP algorithm.

<u>Proof of correctness:</u> Any interesting path in *G* can be at most *n* − 1 edges long. As a particular edge could appear anywhere along such a path, its rank can range between 1 and *n* − 1. Hence the *m* × (*n* − 1) recurrence table *T* sufficiently captures all possibilities for each edge in *E*. The following key observation completes the proof. Let *P*\*(*i, j*) denote an optimal scoring path, if one exists, of length *j* ∈ [1,*n* − 1] ending at edge *e_i_* ∈ *E*. If *P*\*(*i,j*) exists and if *j* > 1, then there should also exist *P*\*(*i*′, *j* − 1) where *i*′ ∈ Pred(*e_i_*). Furthermore, the edge *e_i_ could not* have appeared in *P*\*(*i*′, *j* − 1) because *G* is acyclic. Therefore, due to the edge-disjoint nature of *P*\*(*i*′,*j* − 1) and the remainder of *P*\*(*i, j*) (which is *e_i_*), the principle of optimality is preserved—i.e., the maximum operator in Eqn. (4) is guaranteed to ensure optimality of *T*(*i, j*).

<u>Complexity analysis:</u> The above dynamic programming algorithm can be implemented to run in *O*(*mn*) space and a worst-case time complexity of *O*(*mnd*_in_), where *d*_in_ denotes the maximum indegree of any vertex in *V*.

<u>Algorithmic improvements:</u> The above dynamic programming algorithm for MAX-IP for DAGs can be implemented to run in space and time smaller in practice than the worst case limits suggested above. First, we note that computing the full table *T* is likely to be wasteful, as it is likely to be sparse in practice. The sparsity of *T* follows from the observation that an interesting path of length *j* ending at edge ei can exist only if there exists at least one other interesting path of length *j* − 1 ending at one of *e_i_*’s predecessor edges. We can exploit this property by designing an iterative implementation as follows.

Instead of storing the entire table *T*, we store only the rows (edges), and introduce columns on a “need basis” by maintaining a dynamic list *L*(*e_i_*) of column indices for each edge *e_i_*.

S1) Initially, we assign *L*(*e_i_*) = {1}, as each edge is guaranteed to be in an interesting path of length at least 1 (the path consisting of the edge by itself).
S2) In general, the algorithm performs multiple iterations; within each iteration, we visit and update the dynamic lists for all edges in *E* as follows. For every edge *e*_*i*′_ ∈ Pred(*e_i_*), *L*(*e_i_*) = *L*(*e_i_*) ∪ {*ℓ* + 1|*℘* ∈ *L*(*e*_*i*′_)}. The algorithm iterates until there is no further change in the lists for any of the edges.

The number of iterations in the above implementation can be bounded by the length of the longest path in the DAG (i.e., the diameter *δ*_max_), which is less than *n*. Also, we implement the list update from predecessors to successors such that each edge is visited only a constant number of times (despite the varying products of in- and out-degrees at different vertices). To this end, we implement the update in S2 as a two-step process: first, performing a union of all lists from the predecessor edges of the form (*, *v*) so that the merged lists can be used to update the lists of all the successor edges of the form (*v*, *). Thus the work in each iteration is bounded by *O*(*m*).

Taken together, even in the worst-case scenario of (*δ*_max_ + 1) iterations, the overall time to construct these dynamic lists is *O*(*mδ*_max_). Furthermore, during the list construction process, if one were to carefully store the predecessor locations using pointers, then the computation of the *T*(*i, j*) recurrence in each cell can be executed in time proportional to the number of *non-empty* predecessor values in the table. Overall, this revised algorithm can be implemented to run in time *O*(*mδ*_max_*d*_in_), and in space proportional to the number of non-zero values in the matrix.

Further, the above implementation is also inherently parallel since the list value at an edge in the current iteration depends only on the list values of its predecessors from the previous iteration.

#### 4.2.2 *An Efficient Heuristic for* IP

In addition to an exact algorithm for MAX-IP (Section 4.2.1), we also present an efficient heuristic for finding IP. The IP formulation aims at identifying a *set* of edge-disjoint interesting paths in *G* such that the overall sum of their scores is maximized. IP is relevant in contexts where the user is interested not only in the maximum-scoring path but also multiple others that cover different parts (and hence different subpopulations) of *G*. Once identified, these paths can be rank ordered in descending order of their scores for display purposes.

Algorithm 1 shows the pseudocode for our IP heuristic. The approach is a simple greedy strategy, in which we iteratively find the next best scoring path (by calling MAX-IP), add it to the working set of paths, and remove all edges of that path from the graph. This procedure is carried out until there are no more edges left. The algorithm has a worst-case runtime complexity of *O*(*m*^2^*δ*_max_*d*_in_).

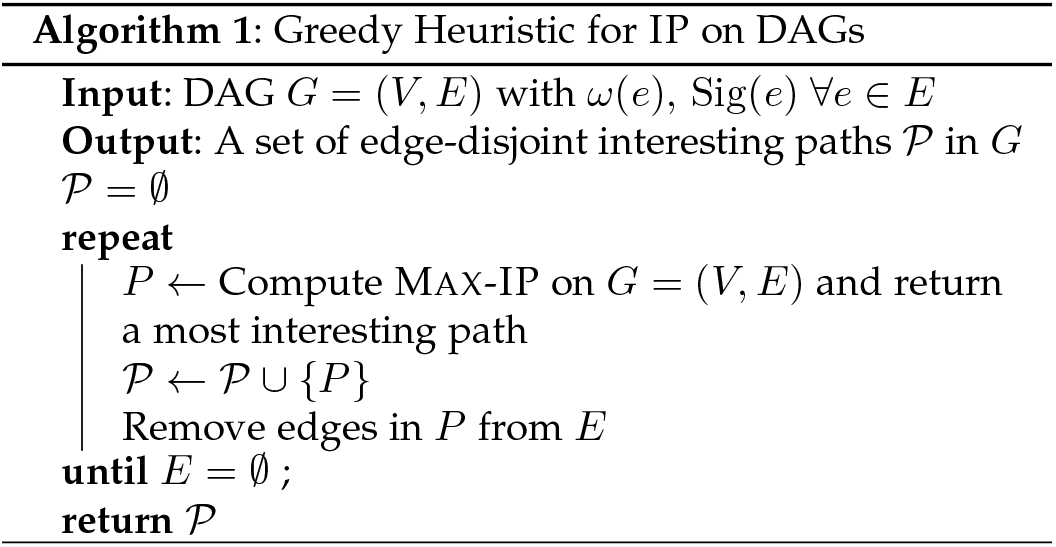

## 5 Experimental evaluation

In our experiments, we used two real-world maize data sets. For the first batch of experiments described in Section 5.1, we used a maize data set containing growth information of two maize genotypes that were cultivated in two different locations in the U.S. (Kansas and Nebraska). We refer to this data set as the “KS/NE” data set. We used this data set to test various functionalities of our `Hyppo-X` framework including hypotheses extraction in the case of single filter function (Section 5.2) and two filter functions (Section 5.3) using both flares and paths. For the second batch of experiments, we used another maize data set collected from two field locations in Nebraska that had identical conditions except for one environmental parameter—one location was irrigated while the other was not. This data set covers individuals from 80 genotypes (as described in Section 5.4). We refer to this data set as “irrigation-controlled” data set.

### 5.1 KS/NE data set

This maize data set consists of phenotypic and environmental measurements for two genotypes (abbreviated here for simplicity as *A* and *B*), grown in two geographic locations (Nebraska (NE) and Kansas (KS)). The data consists of *daily* measurements of the genotypes’ growth rate alongside multiple environmental variables, over the course of the first 100 days of the growing season. For the purpose of our analysis we treat each unique [genotype, location, time] combination as a “point”. Consequently, the above data set consists of *N* = 400 points. Here, “time” is measured in Days After Planting (DAP). An “individual” in this data set refers to a plant individual that corresponds to a [genotype, location] combination. Each point has one phenotypic value (observed growth rate) and 10 environmental variables, including (among others) humidity, temperature, rainfall, solar radiation, soil moisture, and soil temperature.

To study flares and paths, we constructed topological objects out of the KS/NE data set, using single and two filter function(s).

### 5.2 Single filter function

First, we constructed our topological object using DAP as the filter and used the difference in growth rates to calculate pairwise distances between points (in the clustering step). This study is aimed at understanding how the population of individuals (of both genotypes in both locations) show varying trends in phenotypic performance (i.e., growth rate here) as a function of time.

The resulting object along with the detected flares are shown in Fig. 9, based on which we make the following observations.

1. Until around DAP ~40, all four subpopulations behave similarly (as shown by the leading trail of clusters).
2. Around DAP ~40, two branching events emerge: i) The first branching event occurs when the {KS,B} subpopulation separates from the rest due to a significantly accelerated growth spurt (compared to the rest). ii) The second event corresponds to the {KS,A} subpopulation separating from the rest. Fig. 9(B) shows the cluster nodes colored by growth rate.
3. It is not until DAP ~70 that the Nebraska varieties show a similar separation in their behavior.

**Fig. 9.**
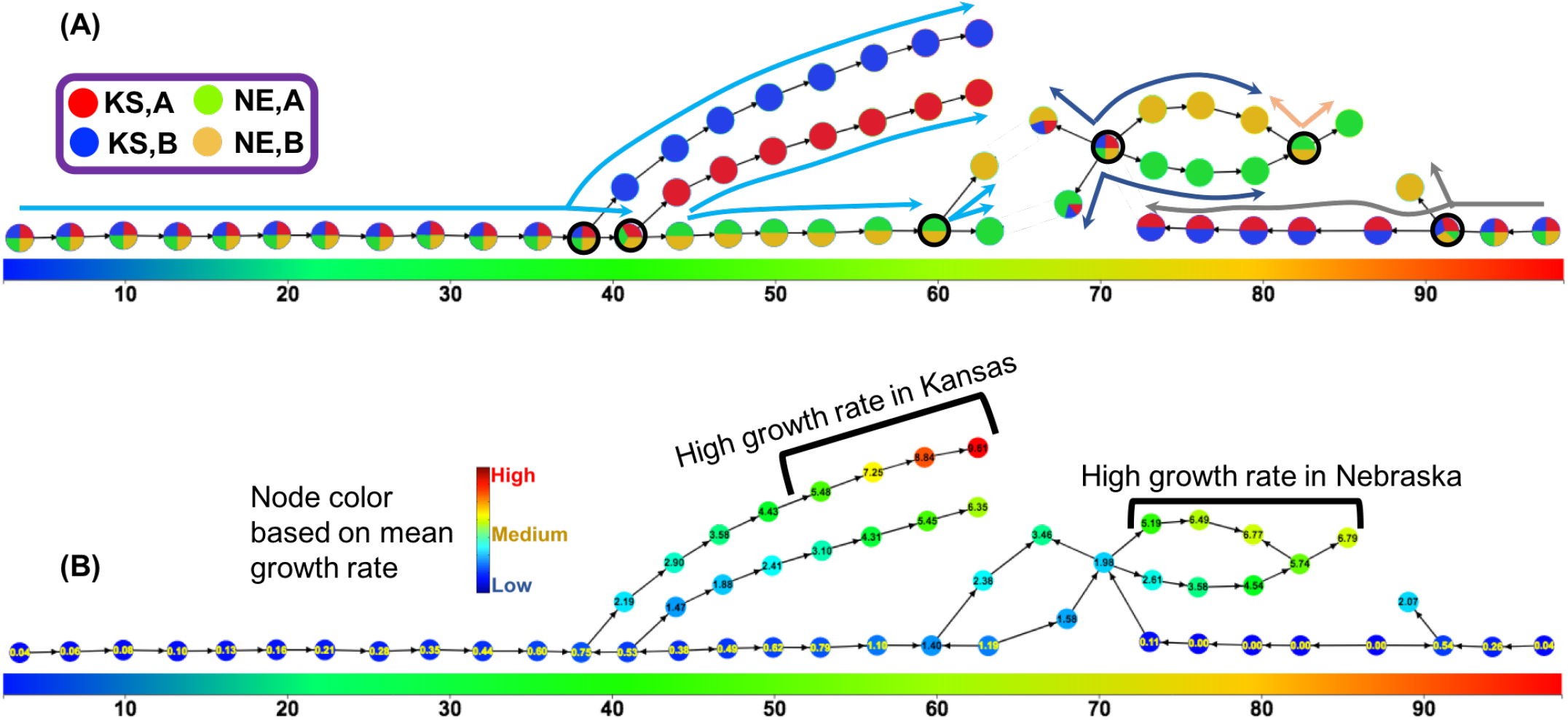
Topological object constructed using DAP as a single filter function and our method detected interesting flares from the object. The horizontal color bar indicates the gradient of DAP, with the value increasing from left to right. (A) Each cluster (node) of the topological object is rendered as a pie-chart showing the distribution of their four classes of individuals. Long arcs of different colors show interesting flares, and the corresponding branching nodes are identified with bold border. The blue flare (long arc spanning DAP 1 through 60) was ranked as the top interesting flare. (B) Each cluster colored by its mean growth rate (phenotype), with branches showing active growth (high phenotype) marked.

All the above branching events were successfully detected by our flare detection algorithm (shown by long arcs of different colors) in a runtime of 9 milliseconds after the Mapper graph is built. The runtime to construct mapper graph from the KS/NE data set was 167 milliseconds. Note that our method is *unsupervised*—the information about the source genotypes and locations (pie-chart distribution in Fig. 9(A)) was applied only *after* the analysis was completed, just to aid in our interpretation. These results demonstrate our method’s ability to successfully delineate interesting subpopulations that show divergent behavior in an unsupervised manner.

Our path detection algorithm also identified interesting paths in the object of Fig. 9 which are already covered by either a stem of a flare or a branch of a flare or both. Therefore, the observations we can make based on paths using the single filter function DAP are similar to ones we made using flares.

### 5.3 Two filter functions

In the results for single filter function given above, the fact that genotype B in Kansas shows a significantly altered behavior compared to the same genotype in Nebraska indicates that there could be causal environmental factors at play that influence the phenotype. To better characterize such potential candidates for key environmental variables, we conduct two-filter studies (one filter being time or DAP, and another filter being one of the many environmental variables recorded). We explored choices of a multitude of environmental variables. In the interest of space, we present the results for {DAP, humidity} combination as it led to more interesting observations compared to other variables.

**Flares**: Fig. 10 shows the corresponding topological object on which we show flares. Based on this figure, we make the following observations:

1. Fig. 10(A) shows that in the initial growth period (1-10 DAP), the performance at both locations are highly comparable, as is evidenced by the clustering of both locations.
2. Around DAP 11 the locations diverge into two separate branches (as shown in panel (A)). This separation is correlated with variation in local humidity values (see panel (C))—more specifically, while Nebraska experienced steadily low humidity values until around DAP 50, Kansas experienced fluctuating and often high humid conditions for most of the period until around DAP 60. This period of high humidity fluctuation also coincides with the accelerated growth rate that Kansas experiences from around DAP 40 (panel (B)). As for Nebraska, the increase in growth rates occur eventually around DAP 60 (panel (B)) and that too coincides with higher values in humidity (panel (C)).

**Fig. 10.**
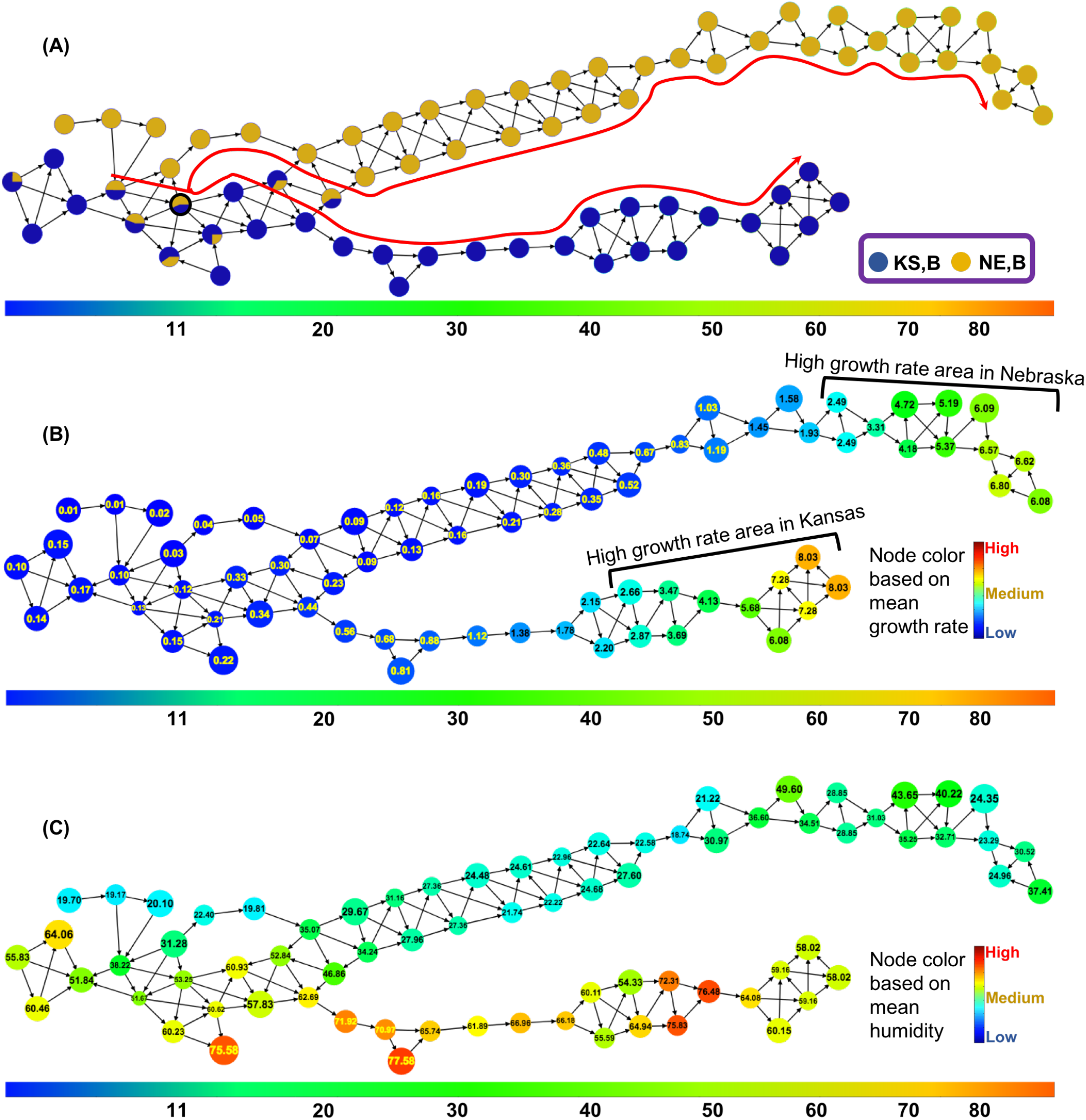
The topological object constructed using only the individuals of genotype B, using DAP and humidity as the two filter functions. The horizontal color bar indicates the gradient of DAP, with its value increasing from left to right. (A) Each cluster (node) is rendered as a pie-chart showing the distribution of its individuals from the two locations (KS and NE) for genotype B. Parts (B) and (C) show the same topological object, however with each cluster (node) colored by the growth rate (phenotype) and humidity (environment), respectively. Our method captured one large flare, which is indicated by the red branched arc in Part (A).

**Path analysis**: In the next step, we ran our interesting path detection algorithm, as described in Section 4.2. All runs were performed with the following settings: i) each path detected is such that all its edges have the same signature; and ii) each path should have at least 3 edges.

The collection of paths that were detected by our algorithm roughly divide the topological object (Fig. 11(B)) into three growth stages at each location (Kansas/Nebraska) based on the growth-rate of plants with respect to the time (DAP). These growth stages are a) Early growth stage, b) Mid-growth stage, and c) Mature growth stage—as shown in Fig. 11:

- **Early growth stage**: The collection of co-located paths *P*_7_, *P*_10_, *P*_11_, *P*_12_ helps us understand how the genotype behaves in its early stages of development in the two locations. More specifically, both paths [*P*_7_, *P*_11_] capture nodes that contain points from both locations because their performances in similar conditions (DAP and humidity) are also quite similar; however, after roughly 22 days after planting (Fig. 11C), the points from KS and NE separate (into *P*_10_ and *P*_12_ respectively).
- **Mid-growth stage in Kansas**: The sequence of paths [*P*_6_,*P*_5_,*P*_1_], which also includes the *most* interesting path by interestingness score (P_1_), represents the active growth period for the KS population (see Fig. 11B). In this period, the growth rate increased from 1.38 cm/day to 8.03 cm/day, from approximately 35 days after planting to 61 days after planting (see Fig. 11C). In contrast, the plants in NE, despite being the same genotype, had very low growth rates during roughly the same period in time (39 days after planting to 60 days after planting; see paths [*P*_9_, *P*_8_] of both Figs. 11B and 11C). Incidentally, examining the humidity trends in the same period for these two locations (see Fig. 11D), we see that the humidity was very low in NE compared to KS, and that the increase in humidity values for the NE population (after 56 days after planting) coincides with the increased activity in its growth rate (see Figs. 11C, 11D)—thereby giving us an indicator that humidity may have an active role in NE, perhaps more so than in KS, in accelerating growth rate during the mid-stages of development.
- **Mid-growth stage in Nebraska**: The sequences of paths [*P*_9_, *P*_8_] and [*P*_3_, *P*_4_] represent the active growth period of the NE population (more specifically, the growth burst starts from the middle of the path *P*_8_), where the growth rate increases from 1.19 cm/day to 6.57 cm/day (Fig. 11B). This high activity period starts from approximately 56 days after planting and ends roughly at 80 days after planting. As indicated above, this active growth rate coincides with the period having higher humidity for NE.
- **Mature growth stage**: The path *P*_2_ helps us understand how the genotype behaves in its later stages of development in the two locations. More specifically, path *P*_2_ starts with points from both locations because their performances in similar conditions (DAP and humidity) are also quite similar. However, after roughly 92 days after planting (Fig. 11C), plants in both locations do not grow much.
- The path *P*_7_ and *P*_2_ illustrate that plants of the genotype *B* do *not* grow as much before 22 days after planting and after 92 days after planting, respectively.

**Fig. 11.**
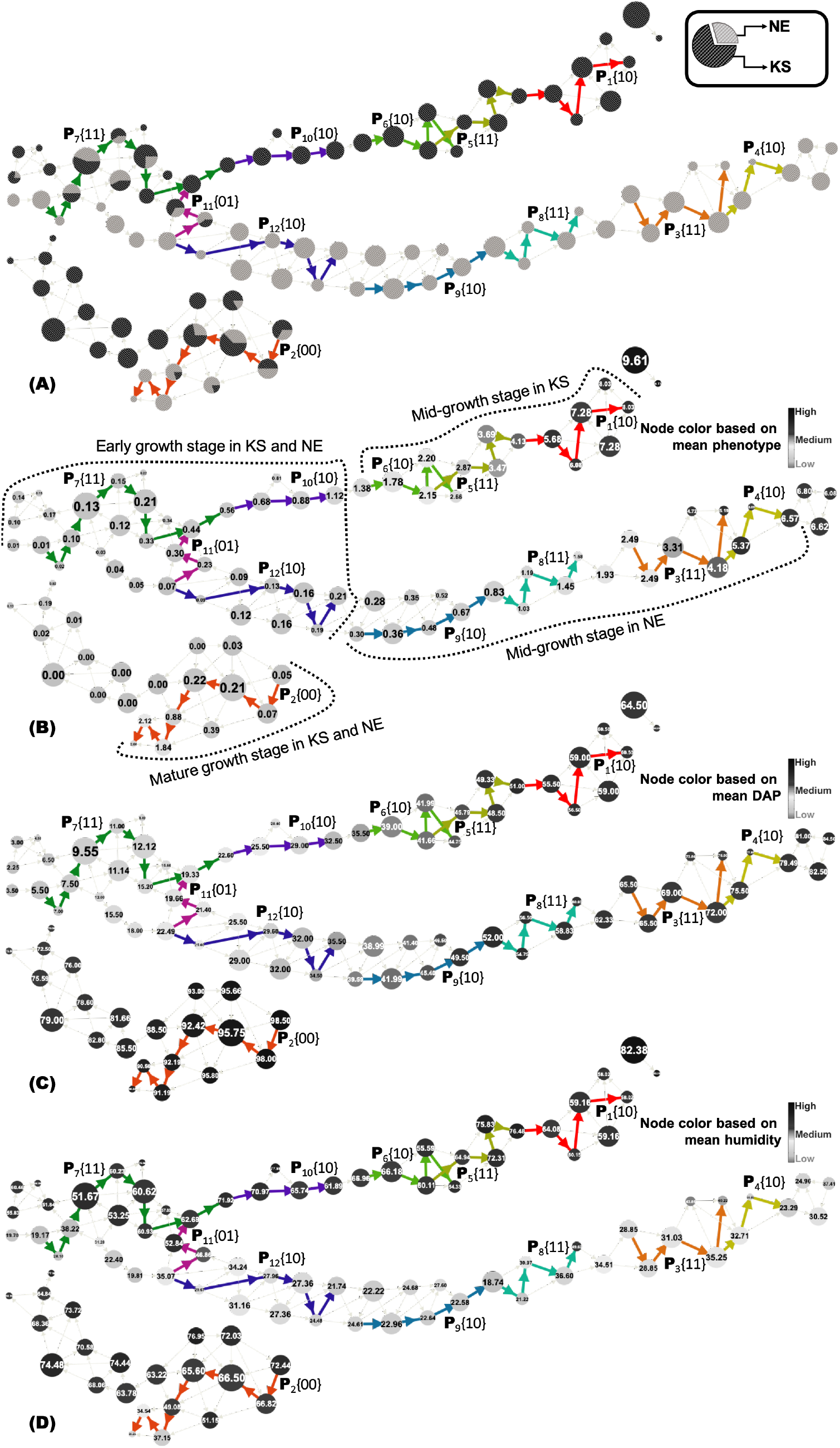
The topological object constructed using DAP and humidity as the two filter functions, using only the [KS,B] and [NE,B] points. The same object is shown in the four panes, albeit with different gray scale coloring schemes. (A) shows each node (i.e., partial cluster) as a pie-chart of the relative concentrations of the two possible locations combinations; (B) shows nodes colored by their mean phenotypic value; (C) shows nodes colored by their mean DAP value; and (D) shows nodes colored by their mean humidity value. The mean values are also indicated within the respective circles. The size of the circle for each node is proportional to the size of the corresponding partial cluster. Also shown highlighted as thick colored edges are the set of interesting paths identified by our method. Edge directions are from low to high mean phenotypic values. The interesting paths are labeled as *P_i_*{*s*_1_*s*_2_}, where *i* is the path number, and *s*_1_*s2* denotes the signature for that path (*s*_1_ corresponds to DAP and *s2* corresponds to humidity). Recall that in the signature, 0 means decreasing and 1 means increasing.

To better understand the results of `Hyppo-X` and contrast the capability of our method with more traditional approaches, we plotted all the genotype B points as a scatter plot, based on their DAP and humidity (see Fig. 1). The coloring of the points are by their location. As can be seen, the plot shows a clear separation between NE and KS humidity values, with NE exposed to lower humidity values than KS, in general. Note that this is a coarse-level information which could have been easily obtained through a correlation test as well. However the limitation of such correlation tests is that they point to global trends without providing insight into the variabilities that may exist across different subpopulations at different scales. On the other hand, identifying such subpopulation-based variability (as output by the paths and flares from `Hyppo-X`) could prove useful in delineating key environmental or temporal triggers that impact crop performance, and on how that behavior varies within a diverse population. That is where our topology-based approach can be useful—to make such inferences from the data and formulate testable hypotheses.

To better illustrate this advantage, we overlaid the interesting path sequences identified by our paths (discussed above) on to the scatter plot. These path sequences are shown as arcs in Fig. 1. As can be seen, our interesting paths show four major “features” within this scatter plot:

i. the initial sequence where both NE and KS varieties behave similarly in their initial developmental stages, before branching out (around 22 DAP);
ii. the period of active growth for [KS,B] between roughly 35 and 61 days after planting; and
iii. the period of active growth for [NE,B] appearing much later, between roughly 56 and 80 days after planting.
iv. The path that separates at around 22 days after planting, merges back after 92 days after planting.

More interestingly, at the end of our interesting paths ([*P*_9_, *P*_8_]) for [NE,B] is also for the first time the humidity value experienced a spike for that location—increasing from values under 35 to around 50*s*—effectively implying (or at least indicating) a probable cause for increased growth activity. After that trigger, minor fluctuations in humidity seemed to have little effect in the growth rate, which continued to increase through 80 days after planting. This study sets up a testable link between a genotype (B) and environmental variable (in this case, humidity) toward a performance trait (growth rate).

These results and observations suggest two plausible hypotheses: (a) that humidity is likely to influence the growth rate; and (b) that this degree of influence is more pronounced on genotype B than for genotype A. The precise time and humidity intervals where such effects manifest are shown by the flare.

This illustrative example serves to demonstrate that our topology-based method also has the potential to enrich further the information that can be obtained through conventional methods such as scatter plots.

### 5.4 Application on Irrigation-controlled data set

#### 5.4.1 Irrigation data set

This maize data set consists of phenotypic and environmental measurements for 80 genotypes, grown in two field locations in Nebraska (NE), USA. Over the growing season, one field location solely depended on rainfall whereas irrigation facility was provided to the other field location. Apart from this irrigation facility, all other environmental parameters are identical in both field locations. The data consists of *daily* measurements of the genotypes’ growth rates alongside multiple environmental variables, over the course of the first 80 days of the growing season. For the purpose of our analysis we treat each unique [genotype, time] combination as a “point”. For each point, we computed the growth rate difference from the irrigated location to the non-irrigated location. Consequently, the above data set consists of *N* = 6400 points. Here, “time” was measured in Days After Planting (DAP). An “individual” in this data set refers to a specific genotypic plant.

**Topological object construction**: First, we constructed our topological object using DAP as a single filter function (parameter setting for this analysis is given in Table 1) and phenotypic difference between points for clustering. This study is aimed at understanding how the population of individuals (genotypes) show varying trends in phenotypic performance (i.e., growth rate here) as a function of time with respect to two distinct controlled environments (irrigated and non-irrigated). The resulting object is shown in Fig. 12.

**Fig. 12.**
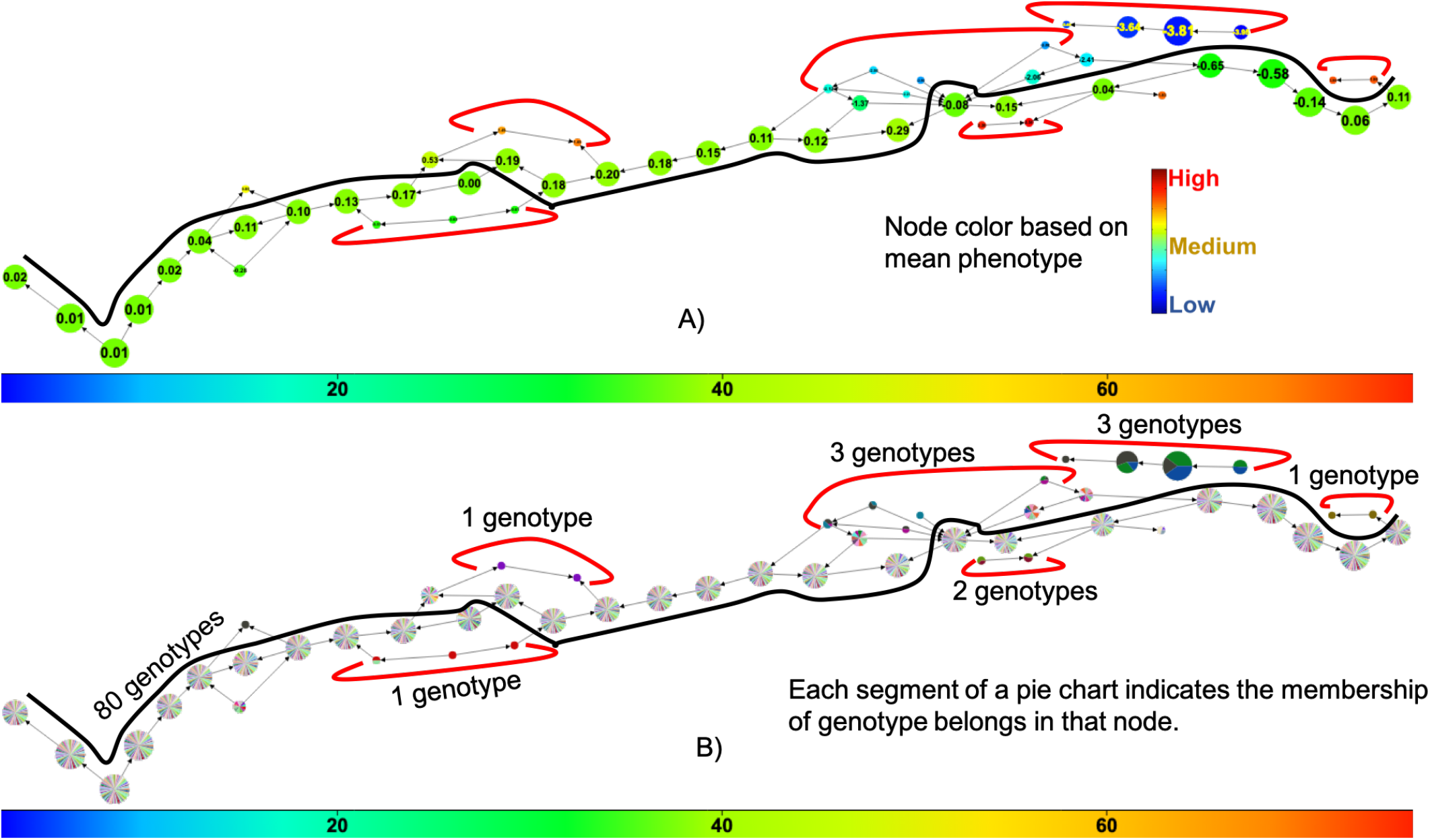
Topological object constructed using DAP as a single filter function. The horizontal color bar indicates the gradient of DAP, with the value increasing from left to right. All the nodes along black colored arc shows almost similar growth rate difference (see part (A)) which means the growth rate difference between irrigated and non-irrigated locations for all the genotypes (see part (B)) that belong in this path are very close to each other. We also observed some nodes (marked by red-colored arcs) with divergent phenotypic values, which indicate that the genotypes (see part (B)) of these nodes have phenotypic variation between two different controlled environments (irrigated and non-irrigated). A detailed overview of these genotypes are given in Appendix A.

**TABLE 1.** Parameter settings for single filter (DAP) analysis on irrigation-controlled dataset.

| Steps | Parameters |
| --- | --- |
| Filtering | 27 windows along filter DAP |
| Clustering | $r = 0.2, \rho = 2$ |
| Persistent overlap | 20% |

**Object exploration:** From the resulting topological object shown in Fig. 12, we observed phenotypic variation of some of the genotypes when they were exposed in two different controlled environments (irrigated and non-irrigated). According to Fig. 12(A), all the nodes along the bold black line show similar performance, which indicates that the points belonging to these nodes have small growth rate variation between irrigated and non-irrigated environments. On the other hand, the nodes those are marked by red-colored arcs contain points which show large growth rate variation between irrigated and non-irrigated environments. The genotypes listed in Table 2 are retrieved from the points belonging to the nodes marked by red arcs (Fig. 12(B)).

**TABLE 2.** List of root worm affected genotypic plants and their corresponding DAP range.

| Genotype | Starting DAP | Ending DAP |
| --- | --- | --- |
| PHW52 x LH123HT | 27 | 36 |
| PHB47 x PHR55 | 33 | 39 |
| LH198 x PHW30 | 54 | 63 |
| PHW52 x Q381 | 51 | 57 |
| PHP02 x PHB47 | 59 | 63 |
| PHB47 x LH185 | 57 | 63 |
| PHB47 x PHG83 | 51 | 67 |
| LH198 x LH51 | 62 | 69 |
| PHB47 x LH38 | 66 | 69 |
| ICI 441 x PHZ51 | 74 | 78 |

The set of genotypes covered in the nodes marked by red arcs are interesting because each of them shows phenotypic variation when exposed to two environments. Originally, our working hypothesis was that this phenotypic divergence is a result of irrigation vs. non-irrigation, or some other factors that affected growth in these plants. Upon careful examination by domain scientists who generated the data (Hey and Schnable, coauthors of this manuscript), we indeed confirmed that the plants selected in these genotypes were affected by a root-worm disease. Root-worm is an insect that cuts roots of a plant, which lead to the death of the plants. Our method is able to detect such affected genotypic plants in a unsupervised way. For a closer look, we zoomed into part of the topological object (Fig. 12(B)) and marked the starting and ending DAP for each of the genotypes listed in Table 2 in Appendix A.

The above application on the irrigation controlled data set also shows the ability of our framework to extract another type of topological feature—one of *spines* where majority of the points follow one behavioral pattern while a small subset of points deviate (as divergent paths). For instance, in Fig. 12 spines are indicated by the thick black arc, while the divergent paths are identified by red arcs. An extension of our framework could be to score and identify interesting spines similar to how we identified flares and paths. Another related extension is one of finding highly traversed paths in the topological object and compare them to less traversed paths.

## 6 Conclusion

We have presented a scalable exploratory framework for navigating high-dimensional data sets and applied it to plant phenomics data to analyze the effect of environmental factors on phenotypic traits. At its core, our approach is fundamentally different from state-of-the-art techniques in many ways as outlined below. First, it inherits the advantages of topology including its use of coordinate-free representations, robustness to noise, and natural rendition of compact representations. Second, by allowing the user to define multiple filter functions, it enables them to study the combined effect of multiple factors on target performance traits. Third, through its clustering and visualization capabilities, it provides a way for domain experts to readily observe emergent behavior among different groups or subpopulations without requiring the knowledge of any priors. This feature enables scientists to identify subpopulations, compare them, and perform more targeted studies to formulate and test hypotheses.

Our approach is scalable in that it can scale to large data sets containing possibly tens of thousands of points, reducing such large data to tens or hundreds of partial clusters, thereby making visualization and exploration possible.

While the scope of this work can be further expanded through application to a broader range of phenomics data collections, the results presented in this paper show a promising application of topology and its role in hypothesis extraction from high-dimensional data sets. Considering the nascency of the phenomics field, tools for users to explore data and help extract plausible hypothesis in a data-guided manner from large-scale complex data will be important going forward.

## Acknowledgments

The research was supported by U.S. National Science Foundation grant DBI 1661348.

## Appendix A Root worm affected genotypes

From the topological object of Fig. 12(B), we visually identified all the genotypes those are listed in Table 2. All the genotypes of this table are affected by root worm disease, that led the affected plants to die. The DAP range in the table indicates a tentative timeframe when plants were affected by this disease. To clearly observe the time duration when a genotype shows performance variation, we zoomed the corresponding portion of the graph. The zoomed portion of the main graph with a short description for each of the genotypes of Table 2 are as follows:

**PHW52 x LH123HT & PHB47 x PHR55**: From Fig. 13 we observed that genotype “PHW52 x LH123HT” shows phenotypic variation from 27 DAP to 36 DAP. Similarly, genotype “PHB47 x PHR55” shows phenotypic variation from 33 DAP to 39 DAP.

**Fig. 13.**
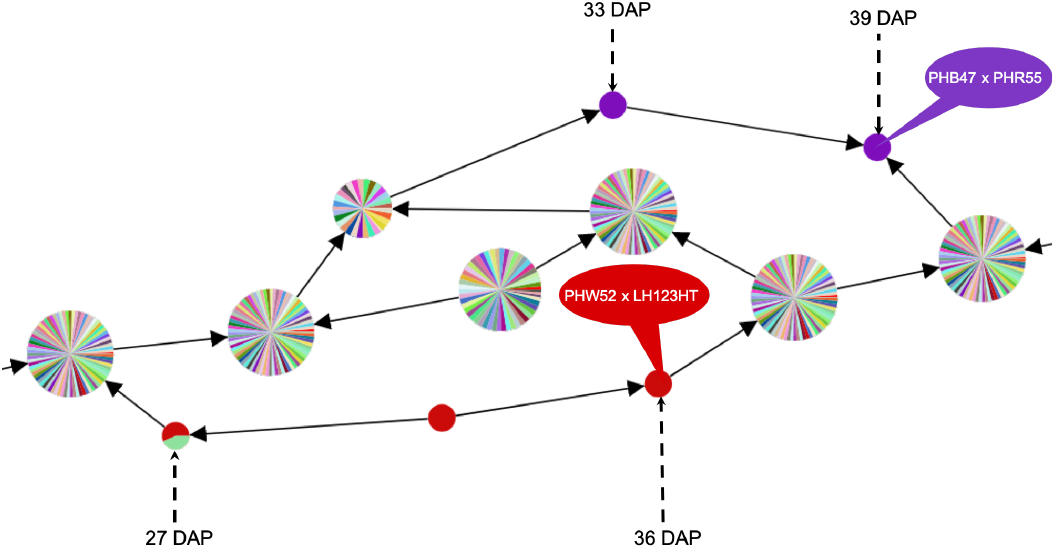
Genotype “PHW52 x LH123HT” shows phenotypic variation from 27 DAP to 36 DAP whereas, genotype “PHB47 x PHR55” shows phenotypic variation from 33 DAP to 39 DAP.

**LH198 x PHW30**: From Fig. 14 we observed that genotype “LH198 x PHW30” shows phenotypic variation from 54 DAP to 63 DAP.

**Fig. 14.**
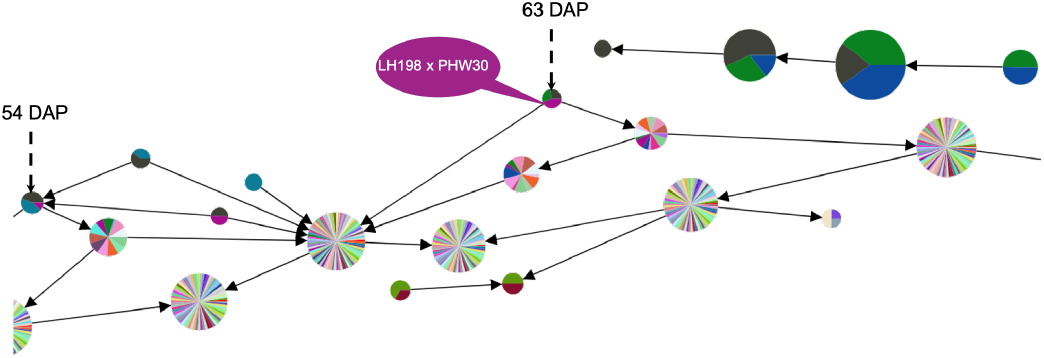
Genotype “LH198 x PHW30” shows phenotypic variation between 54 DAP to 63 DAP.

**PHW52 x Q381**: From Fig. 15 we observed that genotype “PHW52 x Q381” shows phenotypic variation from 51 DAP to 57 DAP.

**Fig. 15.**
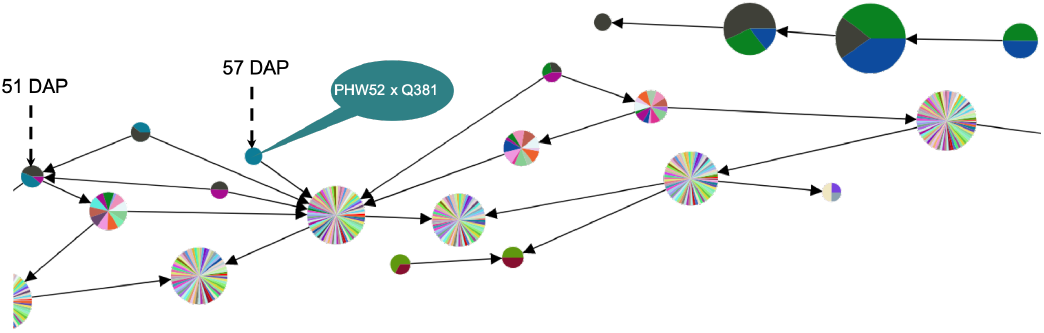
Genotype “PHW52 x Q381” shows phenotypic variation between 51 DAP to 57 DAP.g

**PHP02 x PHB47**: From Fig. 16 we observed that genotype “PHP02 x PHB47” shows phenotypic variation from 59 DAP to 63 DAP.

**Fig. 16.**
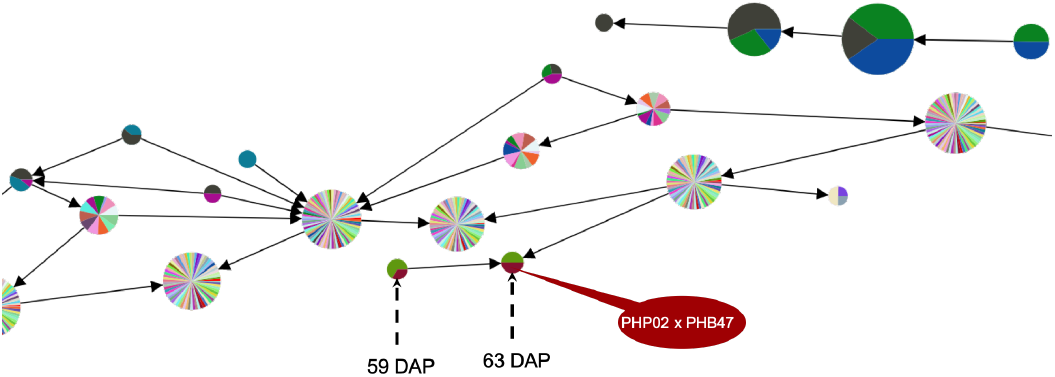
Genotype “PHP02 x PHB47” shows phenotypic variation between 59 DAP to 63 DAP.

**PHB47 x LH185**: From Fig. 17 we observed that genotype “PHB47 x LH185” shows phenotypic variation from 57 DAP to 63 DAP.

**Fig. 17.**
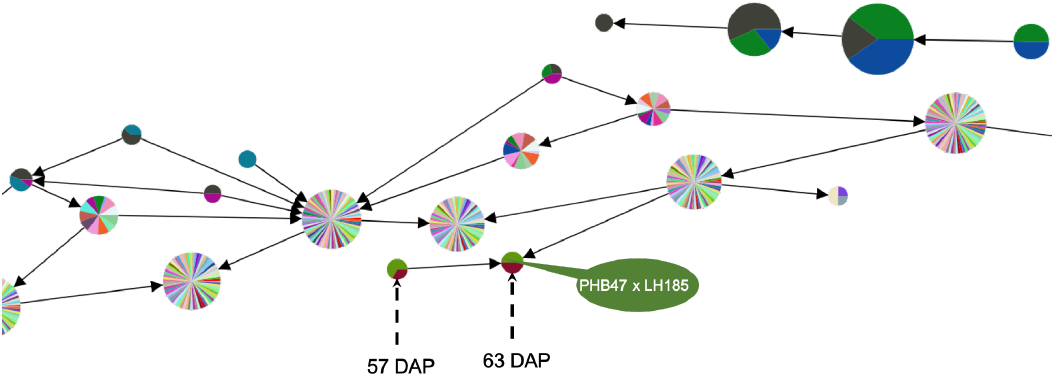
Genotype “PHB47 x LH185” shows phenotypic variation between 57 DAP to 63 DAP.

**PHB47 x PHG83**: From Fig. 18 we observed that genotype “PHB47 x PHG83” shows phenotypic variation from 51 DAP to 67 DAP.

**Fig. 18.**
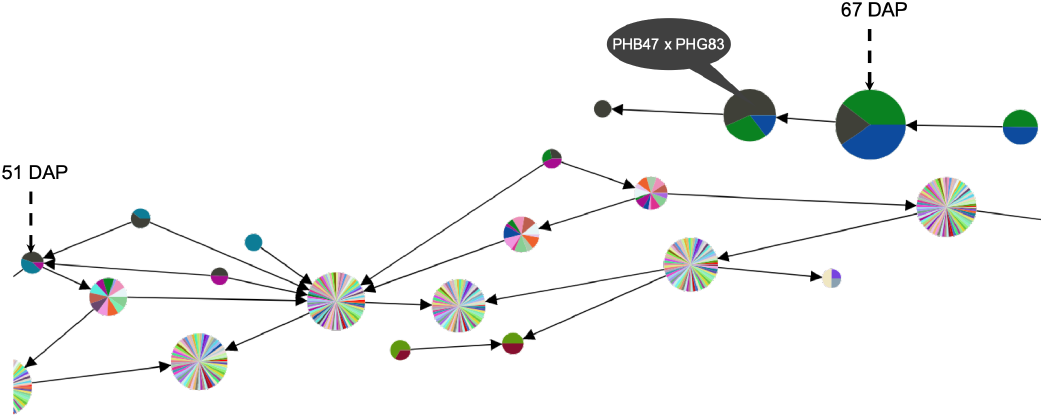
Genotype “PHB47 x PHG83” shows phenotypic variation between 51 DAP to 67 DAP.

**LH198 x LH51**: From Fig. 19 we observed that genotype “LH198 x LH51” shows phenotypic variation from 62 DAP to 69 DAP.

**Fig. 19.**
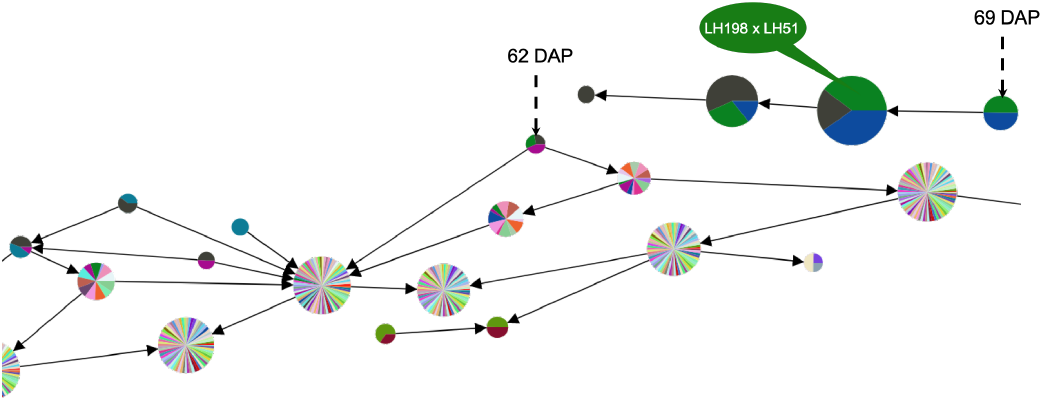
Genotype “LH198 x LH51” shows phenotypic variation between 62 DAP to 69 DAP.

**LPHB47 x LH38**: From Fig. 20 we observed that genotype “PHB47 x LH38” shows phenotypic variation from 66 DAP to 69 DAP.

**Fig. 20.**
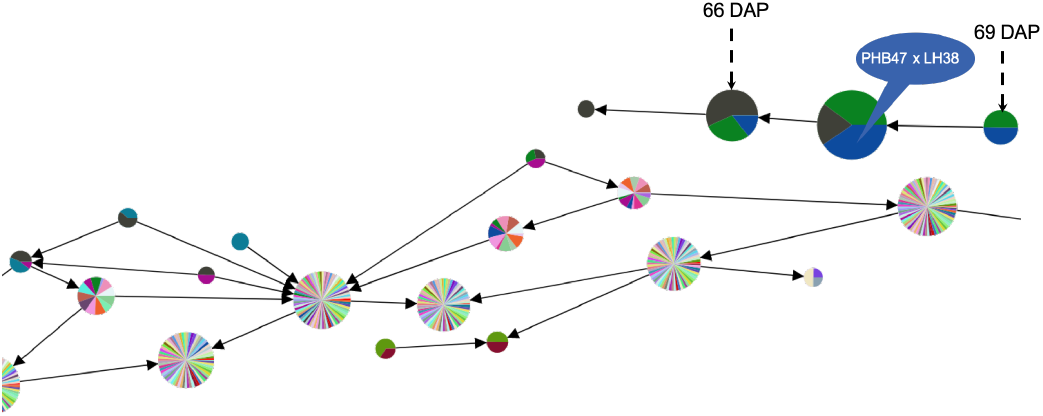
Genotype “PHB47 x LH38” shows phenotypic variation between 66 DAP to 69 DAP.

**ICI 441 x PHZ51**: From Fig. 21 we observed that genotype “ICI 441 x PHZ51” shows phenotypic variation from 74 DAP to 78 DAP.

**Fig. 21.**
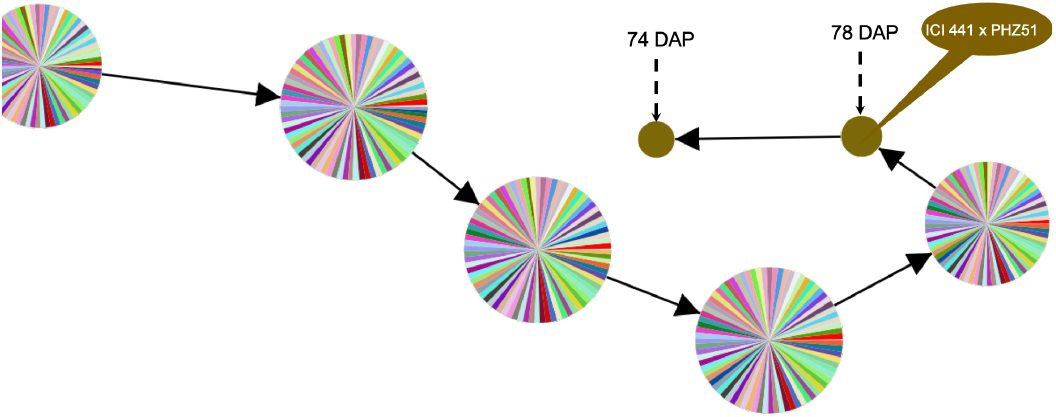
Genotype “ICI 441 x PHZ51” shows phenotypic variation between 74 DAP to 78 DAP.

**Fig. 22.**
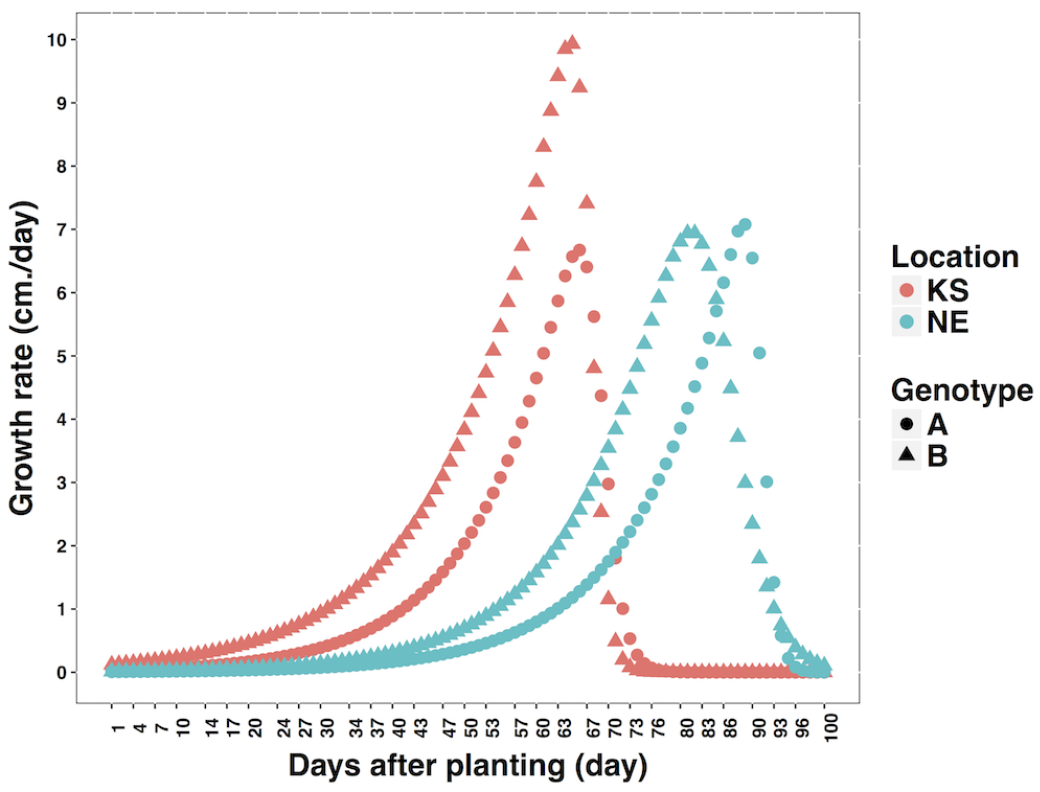
The scatter plot of growth rate with respect to days after planting (DAP). Plants started to grow earlier in location KS compared to location NE. Genotype B of location KS acts differently compared to both genotypes in both locations.

## Appendix B Edge direction in a flare

To capture branches in phenomics data sets accurately, we modify the way in which we direct the edges in the topolog ical object as follows. Given an undirected edge e = {u, v} in the Mapper, we direct edge e from the node with the lower mean phenotypic value to the one with high value, where the respective means are now taken over the subsets of individuals in u and v that belong to genotypes present in both nodes. This procedure is illustrated in Fig. 23.

**Fig. 23.**
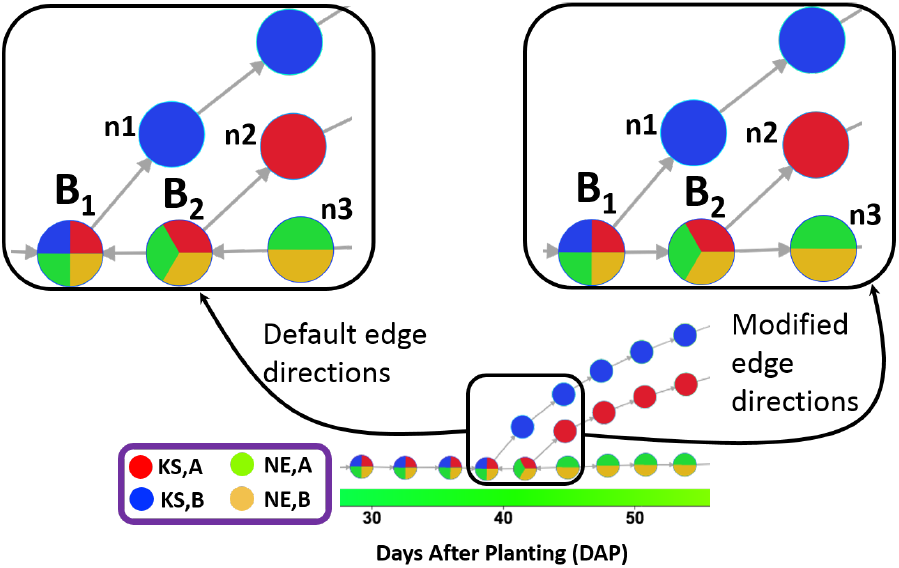
Modification of edge directions in the Mapper. Using the default approach (zoomed in on left), edges are oriented from *B*_2_ to *B*_1_ and from *n*_3_ to *B*_2_, by considering the mean phenotype values of all individuals in these nodes. Using the modified approach (zoomed in on right), we orient the edge from *B*_1_ to *B*_2_ by considering the mean phenotype value of individuals with only the genotypes shared by these nodes (i.e., (KS,A), (NE,A), and (NE,B)). A similar modification directs the edge from *B*_2_ to *n*_3_.

## Appendix C Double filter function (DAP and humidity)

The landscape view of Fig. 24 shows the change of phenotypic value with respect to both time (DAP) and environment (humidity). While some general trends might be evident from this visualization, the question of which subset of points should be compared to which others in order to discern the independent or combined effects of time and environment cannot be answered easily, due to the explosive number of such combinations. In order to delineate the phenotypic variation of a subpopulation under certain environmental condition over a period, we applied topological data analysis methodology. The step by step construction process of our topological object using DAP and humidity as the double filter function are as follows:

**Fig. 24.**
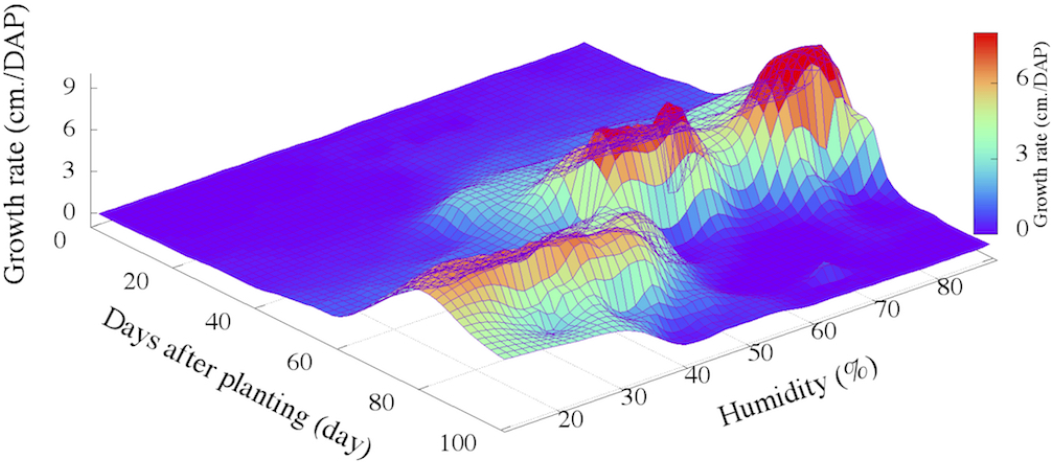
3D representation of phenotype (growth rate) data with respect to time (DAP) and one of the environmental attributes (Humidity) for a real-world maize data set grown in Kansas (KS) and Nebraska (NE). Each point in the landscape is a [genotype, location, date/time] combination.

### C.1 Filtering

The scatter plot of Fig. 1 shows the data points ([genotype, location, date/time]) with respect to two filters, one is time (DAP) and other is environment (humidity). We created 30 windows along the filter DAP and 5 windows along the filter humidity, which creates 150 rectangular interval. Each rectangle has a center point. Initially we did not have any overlapping between two adjacent rectangles. We started from 2.5% overlapping between two adjacent rectangles by increasing the length of each rectangle along both sides from the center point of corresponding rectangle.

### C.2 Partial clusters

In the next step to compute partial clusters, the point set from each rectangular interval was clustered using the algorithm described in Section 3.2. The *distance* between any two points in the set was given by the absolute difference of their trait values. Using the density-based clustering algorithm, we generated a set of partial clusters for every rectangular interval. Every rectangular interval contains a set of points. We calculated standard deviation of phenotypic values for all the points in a rectangle. The mean value of all the standard deviations is used as a clustering radius, which is *r* = 0.7 for this experiment. The density threshold of the clustering is *ρ* =2, which is fixed in our experiments.

### C.3 Simplicial complex

The output of our TDA framework is a simplicial complex constructed using the overlaps among the set of partial clusters generated from all rectangular intervals. Recall that each node represents a partial cluster. In our visual representation of the complex, the size of each node is scaled to its weighted cardinality (as defined by the number of core and peripheral points within that cluster).

### C.4 Persistent homology

As a process to apply homology, which is described in section 3.5, we generated a set of overlapping parameters started from 2.5% to 50% with 2.5% interval. For each overlapping parameter, when a simplicial complex generated, we recorded all the new simplex information against that overlapping value. This overlapping value is considered as a birth or starting point of that new simplex. Similarly, when any existing simplex is replaced by a new simplex (i.e. 1-simplex can be replaced by 2-simplex when a new cluster overlaps two existing clusters of 1-simplex) then that overlapping value is considered as the death or terminating point for the old simplex as well as the birth point for new simplex. After generating simplicial complexes for all the overlapping parameters, we got a list of birth and dead overlapping values of the simplices. This record is used to generate barcode.

Recall from section 3.5, barcode for dimension *zero* indicates the number of connected components and barcode for dimension *one* indicates the number of holes. The barcodes generated from our dataset (Fig. 25) shows that the number of connected components did not change after 42% and the number of holes did not change after 46%. We considered 46% as a constant overlapping value for this double filter function (DAP and Humidity).

**Fig. 25.**
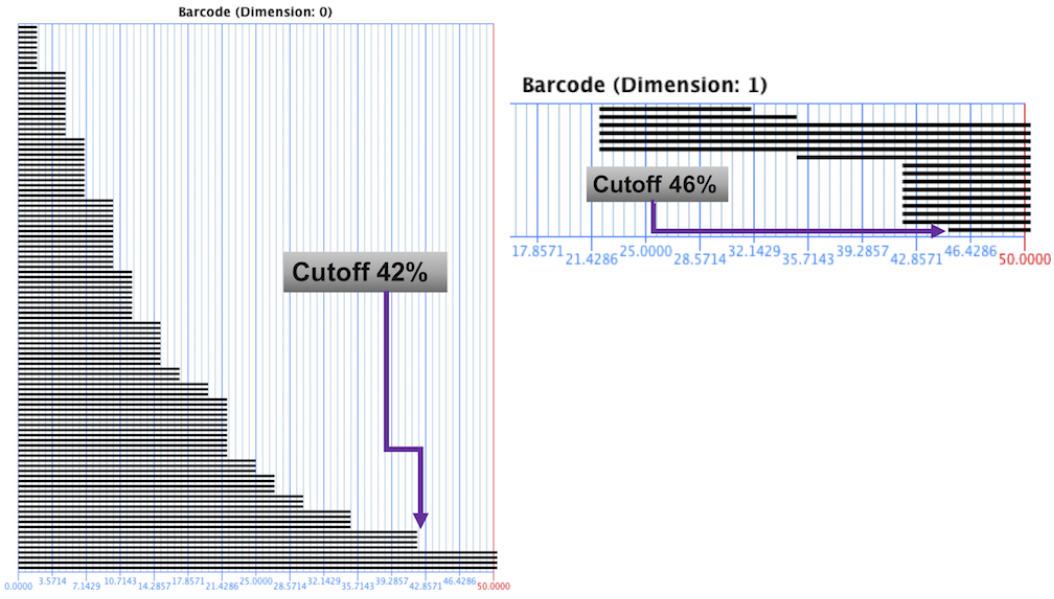
Barcode generated from 0% overlapping to 50% overlapping with 2.5% interval. Barcode of ‘dimension 0’ shows the number of connected components and barcode of ‘dimension 1’ indicates the number of holes. Each horizontal bar specifies the life span of a simplex in terms of percentage overlapping value. The percentage overlapping value after which there has no any terminal point of any simplex is considered as the persistent value. Here, the number of connected components are not changing after 42% and the number of holes are not changing after 46%. These two values are the persistent values and we chose the larger one.

### C.5 Topological object

The topological object in Fig. 11 was generated considering the 46% overlapping. The details analysis of this object is in the result section (Section 5.3).

